# Distinct brain responses to psilocybin and escitalopram in depression captured by the Fluctuation-Dissipation Theorem

**DOI:** 10.64898/2026.06.12.731811

**Authors:** Paulina Clara Dagnino, Irene Acero-Pousa, Gorka Zamora-López, Anira Escrichs, David Erritzoe, David J. Nutt, Robin L. Carhart-Harris, Yonatan Sanz Perl, Morten L. Kringelbach, Gustavo Deco

## Abstract

In recent decades, the psychedelic psilocybin has been studied as a potential treatment for major depressive disorder (MDD), offering an alternative to traditional antidepressants. However, the brain changes underlying the clinical effects of different interventions remain unclear. Here, we investigated the effects of psilocybin and a conventional antidepressant, escitalopram, from the double-blind randomised controlled trial (DB-RCT) -NCT03429075- on the brain’s hierarchical organisation. Using pre- and post-treatment resting-state functional magnetic resonance imaging (fMRI) we built whole-brain models and obtained a generative effective connectivity (GEC) matrix for each patient. Based on the GEC, we measured the level of non-equilibrium brain dynamics by quantifying the deviation from the fluctuation-dissipation theorem (FDT) and performed complementary analysis on brain segregation and asymmetry. Our results showed opposite reconfigurations of the hierarchical non-equilibrium brain dynamics following each treatment. Additionally, baseline measures effectively distinguished responders from non-responders within each treatment. These findings suggest that the deviation of the FDT may serve as a marker for differentiating the effects of psilocybin and escitalopram in MDD treatment, overall, contributing to the understanding of therapeutic mechanisms of depression.

## Introduction

A primary contributor to world-wide disability is major depressive disorder (MDD) (WHO, 2017), significantly impairing quality of life and ordinary everyday activities (Vigo, 2016). Depression is a neuropsychiatric disorder with heterogeneous symptoms, including rumination, negative thoughts, low mood and decreased cognitive flexibility (Beck, 1998). Existing treatments such as selective serotonin reuptake inhibitors (SSRIs) (e.g., escitalopram) present modest efficacy (Hofmann et al., 2017), poor adherence and high relapse rates (Steinert et al., 2014), and negative side effects (Locher et al., 2017). Overall, the global prevalence of MDD and the shortcomings of existing therapies call the need for novel treatments (Holtzheimer & Mayberg, 2011). Among alternative approaches is the psychedelic psilocybin (Nutt & Carhart-Harris, 2021), which has been of scientific interest since the 1950s, halting in 1970 and resuming in the early 1990s (Nichols, 2020). Over recent decades, multiple clinical trials have assessed its potential to alleviate depressive symptoms (Andersen et al., 2021). Ongoing research continues to investigate therapy with psilocybin and classical treatments, focusing on protocol optimisation, efficacy, and the associated changes in complex brain dynamics.

Research on psilocybin using functional magnetic resonance imaging (fMRI) has primarily focused on the acute phase of action, and in healthy individuals, leaving expectations for the unknown time-dependent carry-over evolution both in health and disease. Brain dynamics during the acute phase of a psychedelic experience in health are characteristically dysregulated and unconstrained. This change includes increases in brain entropy (Carhart-Harris, 2025; Girn et al., 2022; Singleton et al., 2022), repertoire of functional connectivity states (Varley et al., 2020; Atasoy et al., 2018; Tagliazucchi, 2014), information flow (Cruzat et al., 2022) and desynchronisation (Siegel et al., 2024). As a treatment for MDD specifically, post-acute effects of psilocybin present increased global integration (Daws et al., 2022) and decreased directedness of the brain’s hierarchy (Deco, 2024). The investigation of the changes in brain dynamics through time with different interventions for MDD, namely psilocybin and escitalopram, is still progressing and evolving.

Interestingly, an emerging field in neuroscience has begun to uncover insights into brain dynamics departing equilibrium from the viewpoint of thermodynamics. Distinct fingerprints of hierarchical organisation and orchestration of the brain are being revealed (Kringelbach et al., 2024). In systems in equilibrium, state transitions maintain a ‘detailed balance’, in the sense that for any two states A and B, the flow of probability from A to B is equal to the flow from B to A. Contrarily, systems in non-equilibrium break this balance, introducing directionality in the flow (i.e., ‘arrow of time’) and producing thermodynamic entropy (Battle et al., 2016; Gnesotto et al., 2018), a key feature of living systems (Schrödinger, 1944). Non-equilibrium dynamics are proving fundamental to brain function, with studies showing that reduced states of consciousness such as coma, sleep and anaesthesia, are closer to equilibrium (García Guzman et al., 2023; Sanz Perl et al., 2021), while cognitive tasks push the brain towards non-equilibrium (Geli, 2025; Lynn et al., 2021). To our knowledge, the studies of (Deco, 2024) and (Socoro-Garrigosa et al., 2025) are the only ones attempting to fully quantify the whole-brain changes in MDD patients after psilocybin and escitalopram treatments from a thermodynamic perspective. The associated non-equilibrium dynamics are yet to be fully understood.

In the brain, non-equilibrium dynamics arise from the asymmetrical relation of feed-forward and feed-backward interactions between regions, which underpin the brain’s functional hierarchy. Measuring this asymmetry is not trivial and a promising framework to do so is the fluctuation dissipation theorem (FDT) (Deco et al., 2023). The FDT establishes a relation between the fluctuations of a system in equilibrium and its response to a small external perturbation. When the system is in non-equilibrium, this relation is disrupted and there is a deviation from the FDT. Therefore, by perturbing a whole-brain model we can quantify the distance to equilibrium through violations of the FDT. Overall, brain asymmetries in information flow cause deviations from the FDT, as such the proportion of the asymmetries can be quantified by the FDT violations, and in turn this deviation from the FDT provides a framework to characterise brain states in terms of hierarchical non-equilibrium dynamics. This approach has been successfully applied to different brain states, showing higher FDT violations (i.e., more non-equilibrium) in wakefulness compared to deep sleep, and in cognitive tasks compared to rest, attributed to elevated computational demands (Deco et al., 2023). On the other hand, lower FDT violations (i.e., less non-equilibrium) in disorders of consciousness compared to healthy controls were identified as a signature of conscious capacity (Martinez-Marin et al., 2025). Finally, higher FDT violations in schizophrenia compared to health were said to reflect stronger non-equilibrium brain dynamics characteristic of the disease (Acero-Pousa Irene, 2024).

The aim of this article is to investigate the hierarchical brain dynamics behind the effects of psilocybin and escitalopram in major depressive disorder. To do so, we implemented the thermodynamic-inspired framework for studying violations of the Fluctuation-Dissipation Theorem in the brain (Deco et al., 2023). We focused on a double-blind randomised controlled trial (DB-RCT) involving psilocybin and escitalopram therapy for MDD with pre- and post-treatment fMRI scans (Carhart-Harris et al., 2021). We built individualised whole-brain models and artificially perturbed them to quantify the FDT deviation for each patient before and after treatment. We hypothesised that each treatment (psilocybin and escitalopram) is characterised by distinct changes in FDT deviations. By analysing these violations of the FDT, alongside metrics of segregation and asymmetry, we successfully demonstrated the significant, opposing brain changes underlying the interventions. In addition, we showed the power of these brain markers in relation to clinical responses. Together, these results contribute to the knowledge and potentially optimisation of treatment of depression with psilocybin and escitalopram.

## Materials and Methods

### Trial

The design of the trial and the primary clinical outcomes has been previously published (clinicaltrials.gov: NCT03429075) (Carhart-Harris et al., 2021; Daws et al., 2022) **(Figure 1A)**. It was conducted at the National Institute for Health Research Imperial Clinical Research Facility and received sponsorship from Imperial College London. It obtained ethical approval (ID 17/LO/0389) from the NHS Research and Imperial College Joint Research and Compliance Office. It also received approval from the Health Research Authority and Medicines and Healthcare Products Regulatory Agency. The study was carried out under a Schedule 1 Drug Licence granted by the UK Home Office. Participants did not receive financial compensation and provided written informed consent.

**Figure 1:**
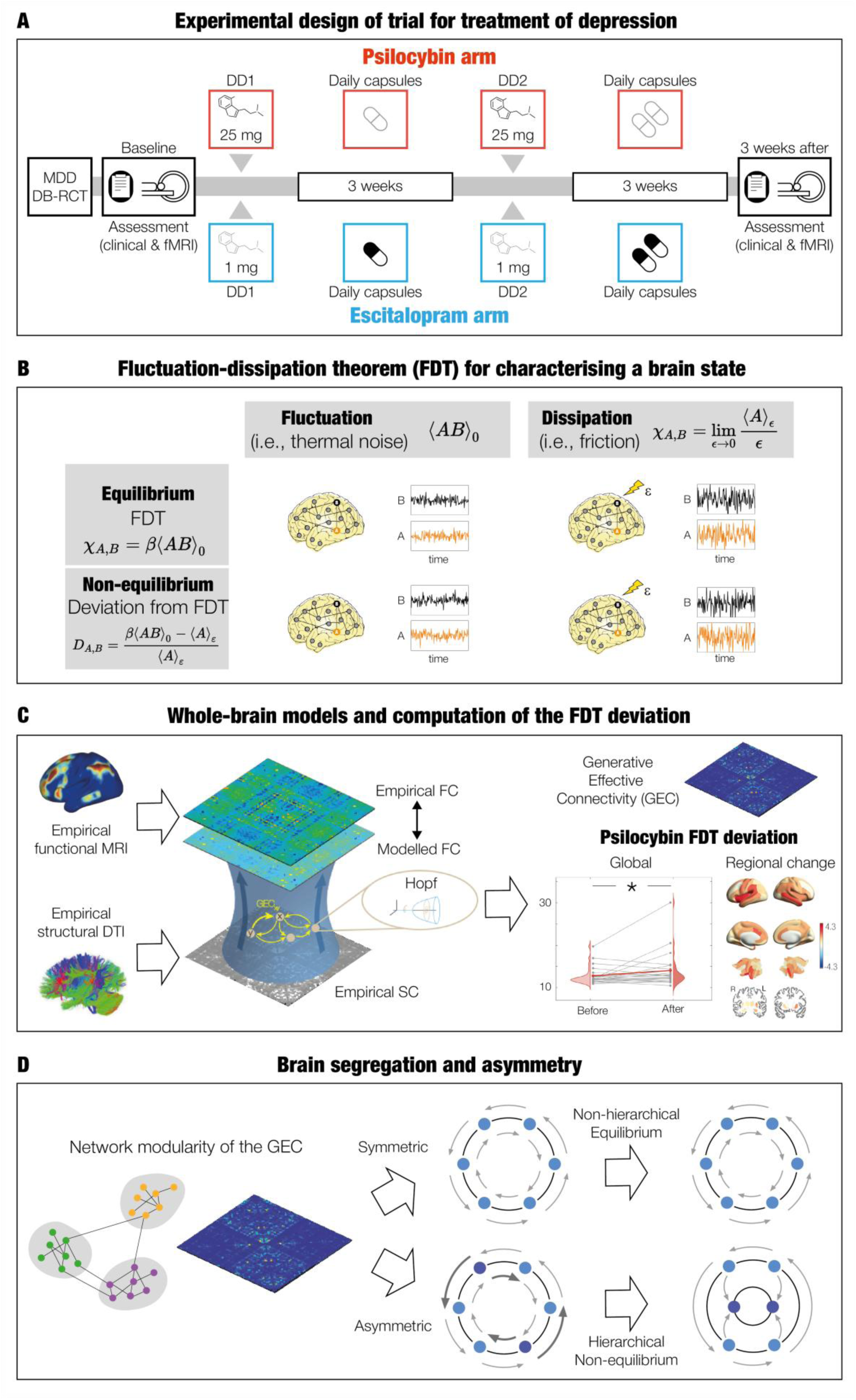
Methodology for capturing differences in brain states before and after different interventions for depression. **A**. Double-blind phase II randomised controlled trial of psilocybin compared to escitalopram (i.e., selective serotonin reuptake inhibitor) treatment. Baseline measures consisted of a clinical assessment and a resting-state functional magnetic resonance imaging acquisition. Patients were randomly assigned to either arm (psilocybin N=22 or escitalopram N=20). Psilocybin treatment involved 2 x 25 mg dosing days (DDs) and 3 weeks of daily placebo capsules after each DD. Escitalopram treatment involved 2 x 1 mg DDs, with 3 weeks of 10 mg and 20 mg and daily escitalopram after DD1 and DD2, respectively. In both arms, patients had a post-treatment clinical assessment and one fMRI scan one day and 3 weeks after DD2. **B.** Fluctuation-dissipation theorem (FDT) applied to empirical neuroimaging data for characterising different brain states. A violation of the FDT establishes that a system is in non-equilibrium, and the level of non-equilibrium can be quantified by the deviation of FDT, in terms of the asymmetric interaction between different brain areas. **C.** Whole-brain models are fitted to empirical neuroimaging data of a given brain state by linking anatomical connectivity with functional connectivity on the basis of generative effective connectivity. These are used to calculate the perturbability maps and the global deviation from FDT. **D.** In the left panel, a brain network is described as a graph consisting of nodes and edges, corresponding to brain areas and their interactions, respectively. Nodes can be divided into communities (i.e., modules), subsets of nodes that have high within-module connectivity and low between module connectivity. A higher quality of partition means a higher extent of separation of non-overlapping modules of brain nodes. In the right panel, asymmetry in the underlying causal interactions of a given brain state can be studied in terms of the hierarchical organisation of the system. The schematic contains two systems with states, represented by circles, and transitions, represented by arrows. A non-hierarchical system (top) has no change in entropy production and is reversible over time. On the other hand, a hierarchical system (bottom) has a change in entropy production and is non-reversible over time due to the asymmetric interactions between different states. Figure adapted from (Deco et al., 2023).

### Participants

Eligibility criteria consisted of practitioner-confirmed diagnosis of unipolar MDD, with a score of 16 or higher on the 21-item Hamilton Depression Rating scale. Individuals were asked if they had prior experience with psychedelics, with 31% and 24% of individuals reporting previous use for the psilocybin and escitalopram groups, respectively. Exclusion criteria were: immediate family or personal history of psychosis, physician-assessed of risky physical health condition, serious suicide attempts history, positive pregnancy test, conditions that prevent undergoing an MRI, contraindications for SSRIs, or previous escitalopram use. Treatment resistance was not an inclusion or exclusion criterion. Eligible individuals had screening interviews via telephone, comprehensive evaluations of mental and physical medical history.

### Treatment protocol

The total number of patients with MDD recruited was 59, from which 30 individuals were allocated to the psilocybin group and 29 individuals were assigned to the escitalopram group using a random number generator. The final samples for this analysis were n=22 in the psilocybin arm (mean age= 41.9 years, s.d. = 11.0, 8 female) and n=20 for in the escitalopram arm (mean age= 38.7 years, s.d. = 11.0, 6 female). The excluded patients for the psilocybin arm consisted in: one patient for choosing not to take daily placebo capsules and COVID-19 UK lockdown, two patients for not attending post-treatment session, five patients from excessive fMRI head motion. The excluded patients for the escitalopram arm consisted in: four patients discontinued from escitalopram’s adverse reactions, one patient with reported cannabis use, one patient lost due to COVID-19 UK lockdown, three patients excluded from excessive fMRI head motion.

Before each treatment, patients had a baseline resting-state fMRI session with eyes closed. The first dosing day (DD1) consisted of either 25 mg of psilocybin, or a presumed negligible dose of 1 mg of psilocybin, for the psilocybin and escitalopram arms, respectively. Participants were informed they received psilocybin but to ensure blinding no information on dosage was provided. Three weeks later there was a second dosing day (DD2), consisting of the same dosage as in the first session for each arm, with no dosage crossover between the two arms. Starting from the first day after DD1, all patients took daily capsules during 6 weeks and 1 day. It consisted of 1 capsule per day during the first 3 weeks, and 2 capsules per day afterwards. In the psilocybin arm, the content of the capsules was inert placebo (microcrystalline cellulose). In the escitalopram arm, the content was 10 mg of escitalopram, resulting in a total of 1 x 10 mg during the first 3 weeks, and 2 x 10 mg afterwards. For blinding, all patients were informed to receive psilocybin but unaware of the dosage. Post-treatment resting-state fMRI with eyes-closed scans were performed for all patients 3 weeks after DD2.

### Treatment outcome

The depression severity assessment used in this study is the Beck Depression Inventory (BDI) - BDI-1A. This measure is patient-rated and captures a wider range of symptoms than other kinds of scoring, with an emphasis in cognitive features of depression (Fried, 2017). The total score range is 0-63, where 0-13 is minimal range, 14-19 is mild, 20-28 is moderate and 29-63 is severe. Assessments were conducted at baseline (before the first dosing day DD1), and at 2, 4 and 6 weeks after DD1. As a note, the measure BDI is the secondary outcome for this study. The primary outcome measure is the Quick Inventory of Depressive Symptomatology (QIDS) – QIDS-SR-16. For justification on the use of BDI and not QIDS please refer to (Weiss et al., 2023).

### Magnetic resonance imaging acquisition

Brain imaging was obtained using a 3T Siemens Tim Trio set-up at Invicro. Anatomical image acquisition was done using the recommended MPRAGE parameters from the Alzheimer’s Disease Neuroimaging Initiative, Grand Opportunity (ADNI-GO56): 1-mm isotropic voxels; 2,300 ms repetition time (TR); 2.98 ms echo time (TE); 160 sagittal slices; 256 x 256 in-plane field of view; 9° flip angle; 240 Hz per pixel bandwidth; 2 generalised autocalibrating partially parallel acquisitions (GRAPPA) acceleration. Functional data (i.e., fMRI) was collected during resting-state for eyes closed, using T2*-weighted echo-planar images and a 32-channel head coil. A total of 480 volumes in ∼10 min were collected with the following parameters: 3-mm isotropic voxels; 1,250 ms TR; 30 ms TE; 44 axial slices; 70° flip angle; 2,232 Hz bandwidth per pixel; 2 GRAPPA acceleration.

### Brain parcellation

Neuroimaging data was parcellated into 80 brain areas using DK80 (Deco et al., 2021). This parcellation combines the Mindboggle-modified Desikan-Killiany parcellation (Desikan et al., 2006) of 62 cortical brain areas (31 areas in each hemisphere) (Klein & Tourville, 2012), with the following 18 subcortical areas (9 areas in each hemisphere): hippocampus, amygdala, subthalamic nucleus (STN), global pallidus internal segment (GPi), global pallidus external segment (GPe), putamen, caudate, nucleus accumbens (NA) and thalamus.

### BOLD fMRI pre-processing

The fMRI data pre-processing was done with a custom in-house pipeline using FMRIB Software Library (FSL) (Smith et al., 2004), Analysis of Functional NeuroImages (AFNI) (Cox, 1996), Freesurfer (Dale et al., 1999) and Advanced Normalization Tools (Avants; Brian B.; Johnson, 2009) packages. Details can be found in (Daws et al., 2022). The stages consisted in de-spiking, slice time correction, motion correction, brain extraction, rigid body registration to anatomical scans, nonlinear template registration, scrubbing, band-pass filtering, regression (with six realignment motion regressors, three tissue signal regressors, draining veins and local white matter). Bias from movement artifacts were ruled out by (Daws et al., 2022).

### Theoretical framework: Violations of the Fluctuation-Dissipation Theorem

Einstein’s work on Brownian motion describes equilibrium systems using the Fluctuation-Dissipation Theorem (FDT), which establishes the balance between friction (i.e., dissipation, transfer of energy) and thermal noise (i.e., spontaneous fluctuations) (Einstein, 1905). Onsager proposed a derivation of the FDT using his regression principle (Crisanti & Ritort; Onsager). This principle states that when a system shifts from an initial equilibrium state towards a final equilibrium state, by a weak external perturbation, this shift can be considered as a spontaneous equilibrium fluctuation (i.e., the initial spontaneous fluctuations predict the dissipation after a perturbation). Conversely, in non-equilibrium systems the spontaneous fluctuations no longer determine the effects of a perturbation (Onsager, 1931).

Let’s assume an observable *B* at time *t* = 0 has a weak external perturbation ε coupled. The expectation value of another observable *A* in the unperturbed state is denoted as ⟨*A*(*t*)⟩_0_, and the expectation value of *A* after the perturbation is applied in *B* is denoted as ⟨*A*(*t*)⟩_ε_. The difference between ⟨*A*(*t*)⟩_ε_ and ⟨*A*(*t*)⟩_0_ is given by:

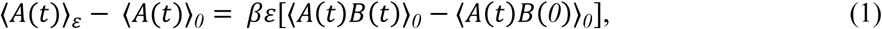

where β is the inverse temperature from equilibrium thermodynamics. Furthermore, the time-dependent susceptibility is as follows:

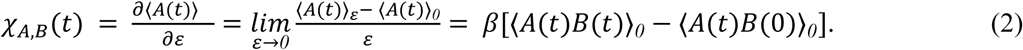

When taking the limit *t* → ∞, the static form of FDT is obtained:

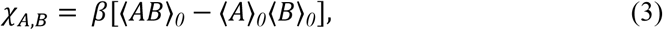

given correlations factorise for infinitely separated times. In equilibrium, we obtain the correspondence between the response of a system to perturbation (the left-hand side of the equation) and the system’s unperturbed correlations (right-sand side of the equation).

By first defining the initial unperturbed state such that the mean values of observables set to zero (i.e., ⟨*A*⟩_0_ = 0 and ⟨*B*⟩_0_ = 0), the level of non-equilibrium can be calculated as the system’s normalised deviation from FDT:

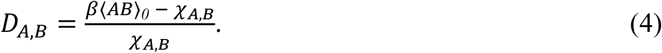

Here, β⟨*AB*⟩_0_ corresponds to the unperturbed fluctuations, and χ_*A*,*B*_ is the response to a small perturbation ε. Finally, the total deviation *D* is calculated by averaging *D*_*A*,*B*_ over all observables *A* and all perturbation sites *B*. In this way, D quantifies the degree of violation of the FDT, hence the distance of the system from equilibrium.

### Model-based FDT of whole-brain data

Following (Deco et al., 2023), we computed the deviation from the FDT of each participant to evaluate the non-equilibrium dynamics in the brain resulting from asymmetries in information flow **(Figure 1B)**. To do so, we built a whole-brain model of each individual fitted to the corresponding functional empirical data **(Figure 1C)**. This allowed us to derive analytical expressions for the correlation between brain areas from the spontaneous fluctuations, and the perturbation effects in each brain region on the average activity across the rest of the brain areas. In other words, we systematically perturb all brain areas *B* and observe the corresponding responses on all brain areas *A*.

### Whole-brain model

The whole-brain model consists of describing the local dynamics of each brain area as the normal form of a supercritical Hopf bifurcation, capable of describing transitions from asynchronous noise to oscillations. From each whole-brain model we created a generative effective connectivity (GEC) matrix. Full mathematical description of the Hopf model, its linearization and optimisation is described in detail in **Supplementary Information**.

### FDT computation

After obtaining a GEC for each participant in each group, we derived an analytical expression for the deviation from the FDT (**Equation 4**). To do so, the expectation values of the state variables ⟨δ*u*⟩_ε*j*_ are calculated when a perturbation ε is applied to a node *j*. From **Supplementary Information Equation 3**, we have the relationship 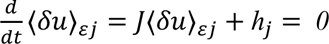, in which *ℎ_j_* is a *2N* vector composed by zeros except component *j* corresponding to the perturbation ε. By solving this for the desired expectation value, the result is ⟨δ*u*⟩_ε*j*_ = −*J*^−^*^1^ℎ*_*j*_. By considering the real part of ⟨δ*u*⟩_ε*j*_, the following is defined: ⟨δ*x*⟩_*j*_ = ⟨δ*x*⟩_ε*j*_/ε. This way, the deviation from the FDT for area *i* when a perturbation is applied to area *j* can be derived as:

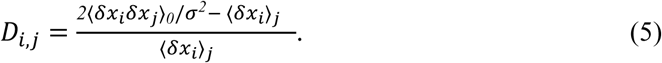

In this equation, the term *2*/σ*^2^*corresponds to the inverse of temperature β and the covariance ⟨δ*x*_*i*_δ*x*⟩*_0_* is derived from *KS^sim^*. Due to numerical motives, the global effect of perturbing node *j* is obtained averaging the numerator and denominator over brain regions:

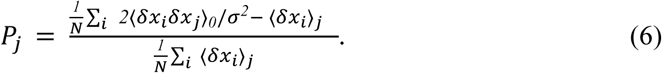

Overall, we obtained for each participant the vector *P*_*j*_, defined as the perturbability map over all brain areas (size *1xN*), corresponding to the effect of perturbing each brain area across the rest of the areas (averaged). In other words, the position of the vector is a perturbed brain area, and the value in the corresponding position is the effect on each of the remaining brain areas at an individual level and then averaged. We finally calculated the level of non-equilibrium 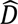 by averaging the deviation from the FDT over all perturbations 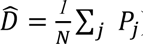.

We also reduced the perturbability map from 80 brain areas to 8 networks: the 7 well-known Yeo networks (Visual, Somatomotor, Dorsal Attention, Ventral Attention, Limbic, Frontoparietal and Default Mode networks) (Yeo et al., 2011), plus the subcortical areas. We used a map of size areas * networks which is computed by counting the number of voxels of each brain area in each network and dividing by the total number of voxels of the corresponding brain area. Thus, each cell contains the probability values of each brain area in each network. For each network, firstly, we multiplied the probability of each area belonging to that network by the area’s perturbability value. Then, we summed the resulting values across all areas and divided by the total sum of probabilities for that network. In other sections of this study, we determine the resting-state network of each brain area by looking at the network with highest associated probability.

### Segregation

To quantify brain segregation (i.e., breakdown of a system into subcomponents), we implemented the Louvain community detection algorithm that assesses the quality of a partition by the modularity index *Q*. This method looks for the maximum extent of separation of non-overlapping modules in a network (Sporns & Betzel, 2016) **(Figure 1D)**.

We implemented the function *community_louvain* from the Brain Connectivity Toolbox in MATLAB, which iteratively moves nodes between communities, aggregates the network, and quantifies the modularity index *Q*. This process is repeated until convergence, calculating the modularity score (i.e., *Q*) 100 times and choosing the highest value as the final result. This output *Q* (i.e., modularity) corresponds to the optimal division of the network into distinct modules (i.e., quality of the network partition) and is considered the measure of segregation. Higher values correspond to a more clearly defined modular organisation (Rubinov & Sporns, 2010).

Modularity is a cost-function that maximises edges within modules and minimises edges across modules. We used Newman as cost-function adapted for weighted networks (Newman & Girvan, 2004). Following (Daws et al., 2022) we first computed the modularity on the FC matrices. Since the modularity algorithm requires positive weights, we applied the absolute value of the FC matrices to retain information from negative correlations. We excluded self-connections by setting the diagonal of the input network to 0, which could otherwise bias the true segregation estimation. For undirected matrices (i.e., FC), *Q* is defined as:

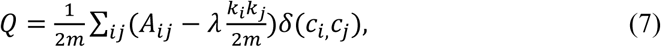

where 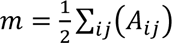. The matrix *A_ij_* corresponds to the weight between node *i* and node *j* of the network and λ is the structural resolution free parameter (set to 1 by default). The term 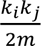 is the expected null network, defined by the total sum of the network across all connections with node *i* as *k*_*i*_ = ∑_*j*_ *A*_*ij*_. In addition, *c*_*i*_ is the community to which a node is assigned to. And, δ(*c*_*i*_, *c*_*j*_) is the Kronecker δ function with a value of 1 if nodes *i* and *j* belong to a same community and a value of 0 if they belong to different communities.

Subsequently, we computed the modularity in the GEC matrices derived from our models. Given the GEC is stored as in vs. out flow of information, we transpose the matrices before computing the modularity in accordance with the computation convention. For directed matrices (i.e., GEC), Q is defined as (Leicht & Newman, 2008):

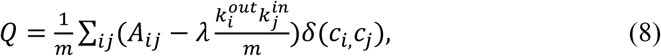

where 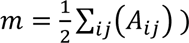. In the implementation, 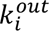 and 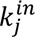 correspond to the outgoing and incoming weights for nodes *i* and *j*.

Lastly, in order to obtain partitions (i.e., modules) representative of each group whilst preserving individual variability, we followed a three-step process from (Bassett et al., 2013). First, we built a nodal association matrix *T* by looking at the number of times each pair of nodes are assigned to a same module in each individual. Then, we divided by the total number of individuals, thus *Tij* indicates the probability of nodes *i* and *j* assigned to the same community. Then, we built a null-model *T^r^* by permutating randomly the initial partitions of each participant whilst preserving the number of modules and module size of the original computation. We set to 0 values of *T* less than the maximum value of *T^r^* to remove noise. Finally, we computed the modularity of the association matrix *T* following the same procedure as described initially (i.e., Louvain algorithm, Newman cost-function, 100 iterations).

### Asymmetry

We assessed the asymmetry in terms of the proportion of asymmetric interactions in a given matrix (global GEC, intra-module GEC and inter-module GEC) **(Figure 1D).** First, we calculated the absolute difference between a matrix and its transpose. Then, we binarised the resulting matrix *A* by converting to zero the cells below a threshold cut-off value γ as follows:

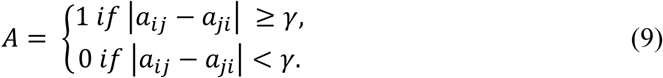

Here, *a*_*ij*_ corresponds to the value of each cell in *A*, and the threshold γ is computed as a percentage of the mean of *A*. Finally, we computed the asymmetry as the proportion of remaining cells (i.e., different to zero) divided by the total number of existing cells.

### Support Vector Machine

We used a support vector machine (SVM) to classify the escitalopram group at baseline into responders and non-responders. The classifier was implemented with the *fitcecoc* function and a Gaussian kernel in MATLAB. We trained the SVM with the leave-one-out cross-validation procedure shuffled 1,000 times, each time randomly excluding one individual and using the rest, with their corresponding class labels, for training. In each iteration, the fully trained two-class model was used to assign a class label to the excluded individual. The final accuracy was the average across all iterations.

We computed a total of 5 classifications using the following different data as inputs: global brain connectivity (GBC), GEC in-weight (GEC^in^), GEC out-weight (GEC^out^), GEC total weight (GEC^total^) and the perturbability map obtained in the main analysis using the FDT framework. The GBC is given by the following equation:

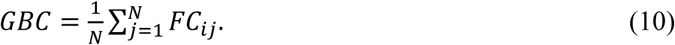

Furthermore, the other calculations on the GEC are as follows:

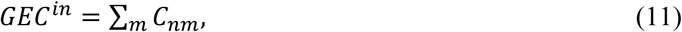

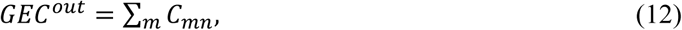

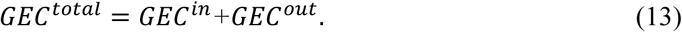

Each input has a size of 1 x *N* (i..e, brain areas). In each of the 5 classifications, we computed the SVM for a number of features *F* (i.e., brain areas) of the input, spanning from 1 to *N* (i.e., 80). Particularly, in each classification (i.e., 5) and number of features *F*, for each k-fold (i.e., 1, 2, 3, …, 1000), we computed the standard error of the mean (SSND) of each brain area between the groups being compared (i.e., responders vs. non-responders) in the training set. Then, we ordered the brain areas in descending order of SSND, and chose the top *F* brain areas. Then, we selected those same brain areas for the testing set.

As a note, the inputs provided by the models (GEC^in^ , GEC^out^ GEC^total^) were calculated for each individual using the DTI as a starting point in the GEC optimisation, not the GEC of the group as for the previous analysis. This way, the goal of the SVM is still to separate underlying patterns between responders and non-responders while lessening the influence of the initial group optimisation.

### Correlation

We studied the relation between baseline brain measures and the level of improvement of the patients by calculating the Pearson correlation between the brain asymmetry within the SOM network at baseline and the change in BDI scores.

### Statistical Analysis

Statistical analysis was performed using permutation-based t-tests with 1000 permutations and a significance threshold of 0.05, paired and non-paired when applicable. For the resting-state network analysis, we applied a False Discovery Rate (FDR) method to correct for multiple comparisons (Hochberg & Benjamini, 1990).

## Results

In this study, we investigated the hierarchical non-equilibrium brain dynamics in a double-blind phase II randomised control trial comparing intervention with psilocybin and escitalopram for MDD patients. Specifically, the fMRI resting-state data at baseline and after treatment **(Figure 1A)** (Carhart-Harris et al., 2021). We used a thermodynamic inspired framework, namely the Fluctuation-Dissipation Theorem (FDT) (Deco et al., 2023), to uncover the changes in non-equilibrium dynamics which are driven by asymmetries between brain areas and that reveal the hierarchical organisation of the whole-brain **(Figure 1B)**. To quantify the FDT in the brain, we created whole-brain models for each participant, fitted to their empirical neuroimaging data. We obtained generative effective connectivity (GEC) matrices, which link anatomical with functional connections through an optimisation process, adjusting anatomical connections with asymmetric weights in an iterative manner. We computed the FDT from the GEC matrices and analysed group differences. Specifically, we perturbed one brain area at a time (e.g., node B) and observed the deviation from the FDT in another area (e.g., node A) (see **Materials and Methods** for detailed explanation). We repeated this for all nodes A after perturbing all nodes B. From this, we constructed a perturbability map in which, for each perturbed area, the deviation of the FDT was averaged over all other areas. We also averaged the perturbability map to obtain the global FDT violation (i.e., deviation from equilibrium) **(Figure 1C)**. In addition, we analysed the GEC by quantifying functional segregation and the proportion of asymmetric interactions (global, intra- and inter- modular) **(Figure 1D)**. Lastly, we evaluated the relation between the brain measures and clinical changes.

### Quantifying the whole-brain FDT deviation in each treatment

At a global level, we found a significant increase of the FDT deviation after psilocybin treatment (p < 0.05), and a significant decrease of the FDT deviation after escitalopram treatment (p < 0.01) **(Figure 2A)**. This reveals a statistically differential treatment-dependent deviation of the FDT. We supported the effect of treatment as the principal factor for the different direction of change using an analysis of variance (ANOVA). The ANOVA analysis showed a significant effect in treatment, and not in the binary classification of responders and non-responders, nor depression scores using the Beck Depression Inventory (BDI) change (after-before) **(Supplementary Information Table S1)**. Patients were classified as responders if they had a decrease in BDI score of 50% from baseline. Furthermore, the Pearson correlation between the change in FDT deviation and the change in BDI scores for each group separately, and together, were not significant (**Supplementary Information Figure S1)**.

**Figure 2:**
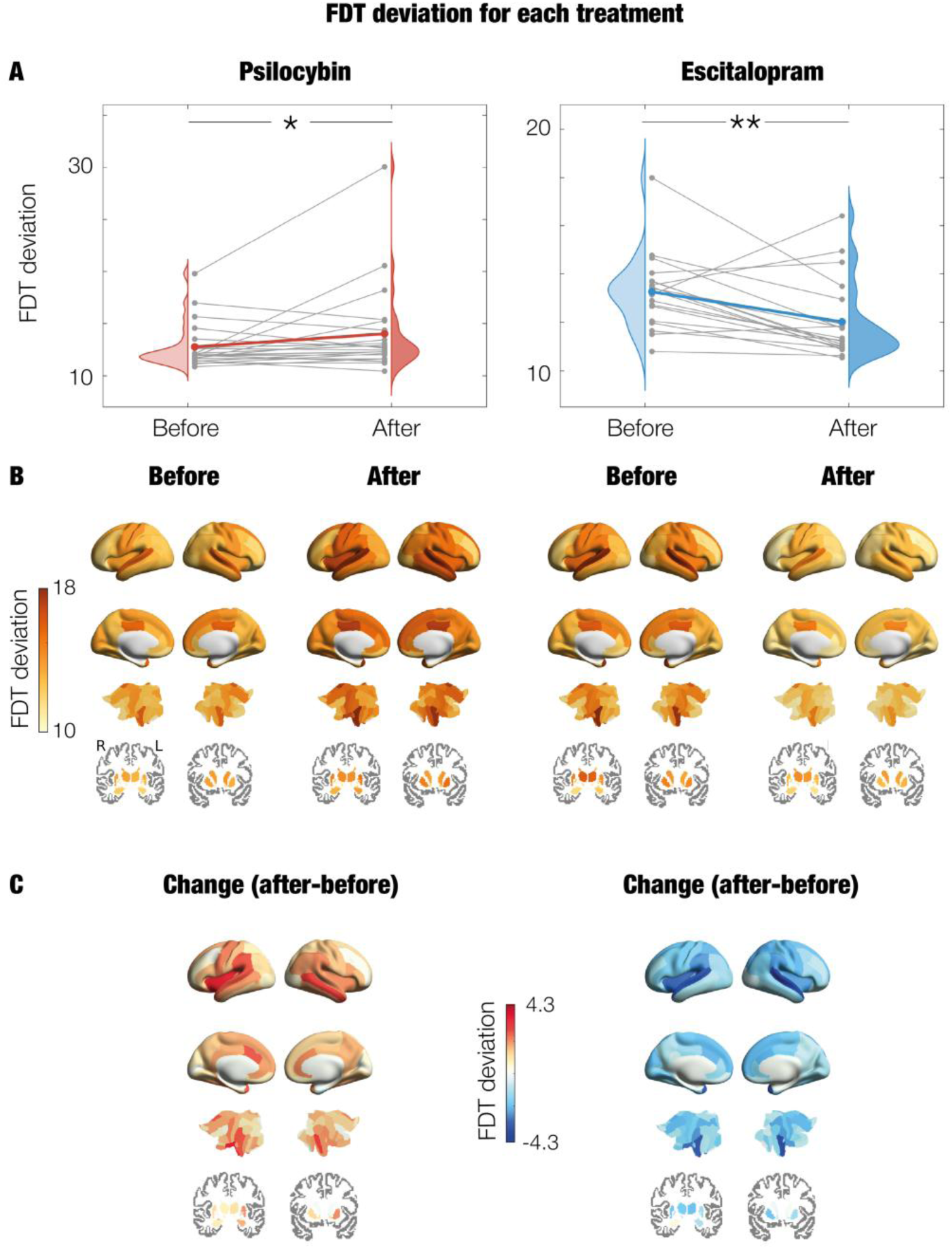
FDT deviation values across the whole-brain following administration of psilocybin and escitalopram. Psilocybin and escitalopram treatments can be observed in the left and right columns, respectively. **A.** Repeated measures plot for the FDT deviation across individuals for each condition. Grey lines represent the trajectory of each individual whereas the coloured line represents their averaged trajectory within each intervention arm (red for psilocybin and blue for escitalopram). There are significant differences between the before and after (* represents p < 0.05; ** represents p < 0.01) for both psilocybin (p = 0.044) and escitalopram (p = 0.004). The change in FDT goes in opposite directions for each treatment, with an increase in psilocybin and a decrease in escitalopram. **B.** Brain renders show the FDT deviation across brain areas (i.e., perturbability map) for the sessions (before and after) of each intervention. Subcortical areas are shown on slices in Montreal Neurological Institute (MNI) space (coronal axis y= −10 and 12 mm). **C.** Brain renders with the difference in FDT deviation (after-before treatment) in each intervention across brain areas. The differential opposite effect for each treatment can be observed.

### Regional differences in FDT deviation

We then carried out an analysis on the FDT deviation at a regional level (**Figure 2B)**. This way, we could evaluate similarities and differences in the patterns of hierarchical reconfiguration between the intervention groups. In both acquisition times (before and after) and treatments (psilocybin and escitalopram), the areas with highest FDT deviation belonged mainly to the somatomotor (SOM) and ventral attention (VAN) networks, as well as some subcortical areas (SCN). Furthermore, the areas with lowest FDT deviation were mainly from the limbic network (LIM), and some areas from the default-mode network (DMN) and SCN **(Supplementary Information Table S2 and S3)**.

With respect to the differences in FDT deviation changes (after-before) within each treatment (**Figure 2C)**, almost all brain areas increased following psilocybin and decreased following escitalopram, in line with their corresponding changes in global FDT deviation. In other words, changes in global FDT deviations are caused by most brain areas within each treatment arm. Under psilocybin treatment, the highest increase was found in areas mainly from the SOM, VAN and DMN. In escitalopram, the largest decrease was found in areas mainly from the SOM, dorsal attention network (DAN), and VAN **(Supplementary Information Table S4)**. Statistics comparing before and after FDT deviation within each treatment revealed less significant brain areas for psilocybin compared to escitalopram (approximately 30 and 50, respectively, with 2 and 37 surviving corrections for multiple comparisons, respectively) **(Supplementary Information Table S5)**.

We also computed the weighted average of FDT deviation across the well-established Yeo resting-state networks (RSN) (Yeo et al., 2011) to evaluate the changes across different spatial scales (global level, regional level, and now RSN level). Psilocybin had an increasing trend of FDT deviation for all networks with significant differences in the SOM, DAN, VAN and DMN (all p < 0.05, non-surviving correction by multiple comparisons) (**Supplementary Information Figure S2A)**. The escitalopram group presented a significant decrease of FDT deviation in all networks: visual network (VIS) and LIM (p < 0.05); SOM, DAN, frontoparietal network (FPN) and DMN (p < 0.01); VAN and SCN (p < 0.001), all surviving correction for multiple comparisons. In both treatments, the SOM and VAN were the networks with highest absolute changes (**Supplementary Information Figure S2B)**. Although the change in these networks was positive for psilocybin and negative for escitalopram, they still had the highest FDT deviation before and after treatment within each group (**Supplementary Information Figure S2C)**.

### Brain segregation

We examined community organisation in the brain network by measuring segregation, the breakdown of a system into subcomponents. We quantified segregation by estimating the modularity, a metric that looks for the highest degree of separation of distinct modules in a network (Sporns & Betzel, 2016). In particular, we employed the Louvain algorithm with Newman’s cost function (Newman & Girvan, 2004). Firstly, to evaluate concordance with the findings of (Daws et al., 2022), we calculated the segregation of functional connectivity (FC) matrices, finding no significance for either group (**Supplementary Information Figure S3)**. We extended the analysis to the GEC matrices. We found a significant difference in modularity after treatment compared to baseline, decreasing for psilocybin (p < 0.001) and increasing for escitalopram (p < 0.001) (**Figure 3A**). This shows the increased power of the GEC compared to the classical FC to differentiate groups and supports the differential opposite statistical effects of psilocybin and escitalopram.

**Figure 3:**
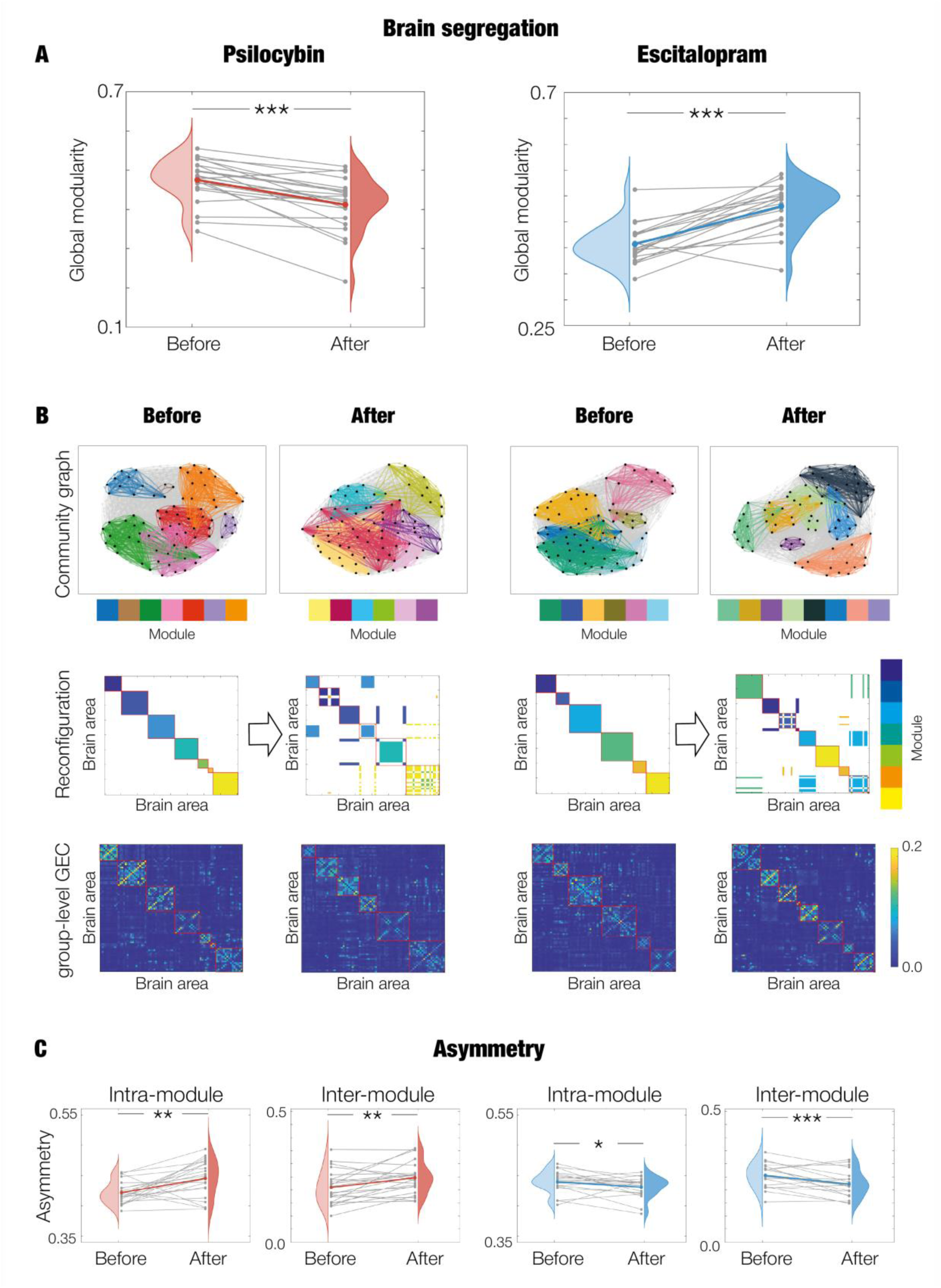
Modularity and asymmetry analysis on the GEC for distinguishing treatment response. **A.** Modularity calculated with Louvain algorithm and Newman as cost-function. Modularity from baseline to post-treatment decreases with psilocybin (p < 0.001) and increases with escitalopram (p < 0.001). **B.** Modularity of the association matrices of each group (i.e., matrix with the probability of two nodes belonging to a same module) following (Bassett et al., 2013) in order to show representative communities whilst preserving inter-subject variability. The first row shows the association matrices with each module in a new colour. Module colours are not comparable between treatments nor before and after within each group. The second row shows the modules coloured according to community affiliation of the baseline within each treatment group. The third row shows the group-level GEC matrices with the modules detected outlined in red. **C.** Intra- and inter- module asymmetry using 90% of the mean of each matrix as threshold, which was the threshold revealing maximum significance in all cases. In all plots, * represents p < 0.05; ** represents p < 0.01 and *** represents p < 0.001. Lastly, grey lines represent the trajectory of each individual whereas the coloured line represents their averaged trajectory within each intervention arm (red for psilocybin and blue for escitalopram).

The correlation between the change in GEC modularity and the change in BDI scores did not give any significance for either treatment arm. When combining both groups (psilocybin and escitalopram), the correlation was significantly positive (**Supplementary Information Figure S4)**. This significant effect after aggregating the groups is masking the fact that treatments show opposite changes in brain organisation (psilocybin leads to a decrease in modularity and escitalopram to an increase in modularity). Therefore, brain modularity changes with treatment, and treatment-specific brain modularity changes are not directly related to BDI changes.

We also obtained representative communities (i.e., modules) of each group by calculating the modularity on the group association matrices (i.e., matrix with the probability of two nodes belonging to a same module) following (Bassett et al., 2013). This method reveals robust modules and preserves variability of individuals. Results in **Figure 3B** show graphs for the association matrix of each group (psilocybin and escitalopram) and session (before and after treatment) with each module in a different colour. After treatment with psilocybin communities are more overlapped, whereas the opposite occurs for escitalopram (i.e., there is more segregation). Furthermore, the reconfiguration of modules from before to after treatment within each group reveals some communities are fully preserved and others are re-organised. Lastly, the number of modules schematised on the average GEC across individuals go in line with the subject-level segregation analysis, decreasing for psilocybin and increasing for escitalopram.

### Brain asymmetry

We then studied the asymmetries in the information flow underlying the deviations from the FDT. We obtained the asymmetry of the GEC matrix by quantifying the proportion of remaining cells after thresholding the absolute difference between the matrix and its transposed (spanning the cut-off value across 10-100% of the mean of the resulting matrix) (**Supplementary Information Figure S5A)**. The global asymmetry significantly increased for psilocybin and decreased for escitalopram, both across all thresholds, in line with the changes in FDT deviation.

Moreover, we calculated the asymmetry in the GEC modules previously found. The intra- and inter-modular asymmetry significantly increased and decreased for psilocybin and escitalopram, respectively (**Figure 3C and Supplementary Figure S5B-C)**. Therefore, within each arm, the change in FDT deviation (and global asymmetric interactions) is driven by both internal and external processing of modules.

### Differences between responders and non-responders

#### Regional differences in FDT deviation

To gain a better understanding of the regional changes underlying the differential hierarchical reconfigurations following response to each treatment, we studied the changes (after-before) in FDT deviation for responders and non-responders (**Figure 4A and Supplementary Information Table S6 and S7)**.

**Figure 4:**
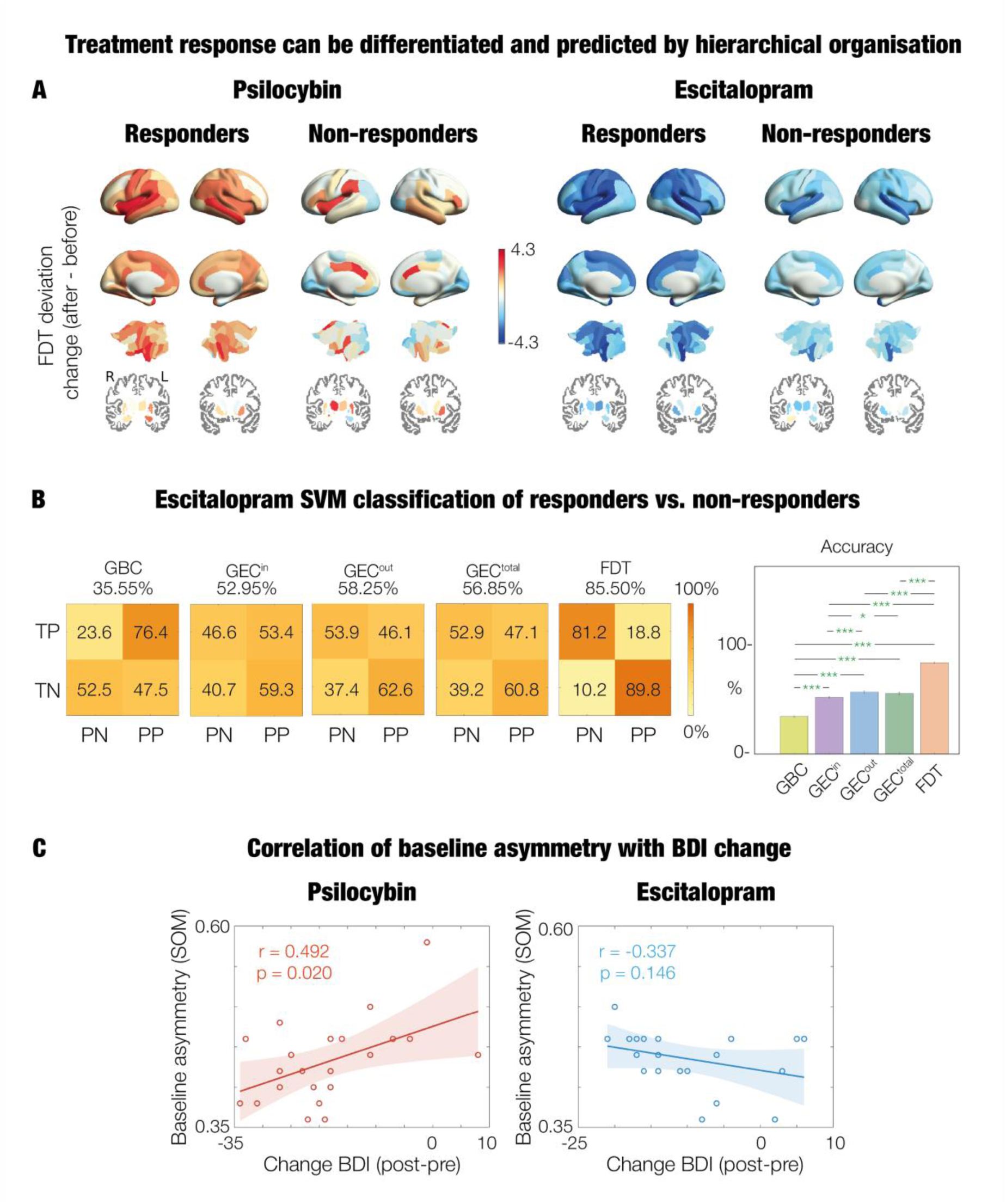
Treatment response can be differentiated and predicted with hierarchical organisation. **A.** Brain renders with the change in FDT deviation in each treatment for responders and non-responders. Subcortical areas are shown on slices in Montreal Neurological Institute (MNI) space (coronal axis y= −10 and 12 mm). **B.** A support vector machine (SVM) was used to predict the treatment response for patients in the escitalopram arm. The analysis could be done given that there are similar number of responders and non-responders for this group, contrarily to psilocybin. We evaluated the performance of the SVM trained with GBC, GEC, and FDT with different number of features (i.e., brain areas) across 1000 k-folds. In the left, confusion matrices show the accuracy as percentages represented in the colour scale, for the maximum accuracy (averaged across 1000 k-folds) reached in each case for a given number of features. TP stands for true positive, TN for true negative, PN for predicted negative and PP for predicted positive. In the right, barplots display the accuracy of each aforementioned SVM, by averaging groups of 10 folds from the total k-folds. The asterisks indicate the significant differences (*, p < 0.05; ***, p < 0.001). The green asterisks correspond to the differences that remain significant after correction by multiple comparisons with FDR. **C.** Correlation of baseline asymmetry within the SOM with the change of BDI scores (after-before). For psilocybin, a significant positive correlation is found, showing that the lower the baseline asymmetry, the higher the improvement in clinical score.

Regarding cortical areas, for psilocybin, responders had positive increases in FDT deviations in almost all brain regions, with the largest positive FDT deviation found mainly in areas associated with the SOM, VAN and DMN. As expected, these changes go in line to the general treatment response, given that most of the patients responded. On the other hand, non-responders had more heterogeneous directions of change in FDT deviation, with half of brain areas changing positively, mainly in the VAN, and the other half negatively, mainly in the VIS and some belonging mainly to the SOM and LIM. For escitalopram, both responder and non-responder groups presented the same direction of change (i.e., negative) in the deviation of FDT, in almost all areas. The responder group showed stronger absolute values in most regions compared to the non-responder group, except some areas from the SCN, as well as VAN and LIM. The highest differences in FDT deviation between before versus after treatment were found for responders in areas mainly from the SOM, VAN and DAN as with the general treatment response, and in non-responders mainly in the VAN.

When focusing on subcortical areas for the responder groups of each treatment, the changes showed different patterns between the intervention arms (**Supplementary Figure S6)**. Firstly, in both treatments, some areas did not follow the respective global trend (i.e., decrease in FDT deviation instead of increase for psilocybin, and increase in FDT deviation instead of decrease for escitalopram). In the psilocybin group, these were the left and right global pallidus internal segment (GPi) and caudate. In the escitalopram group, these were left and right GPi, right hippocampus and left subthalamic nucleus (STN) and nucleus accumbens (NA). Furthermore, with respect to the areas with highest FDT deviation, before treatment these corresponded to the left and right putamen, caudate and thalamus in both psilocybin and escitalopram arms. After treatment, cortical areas replaced these higher positions for psilocybin but not with escitalopram. Regarding the areas with lowest FDT deviations, before treatment these were the left and right amygdala, GPi, STN and NA, both in psilocybin and escitalopram arms. After treatment, cortical areas replaced these lower positions for escitalopram and not in psilocybin. The left and right hippocampus and global pallidus externus (GPe) were maintained in the middle of the FDT deviation distribution across time and treatment, changing in deviation of FDT with the global trend within each group.

### Support Vector Machine for prediction of treatment response in escitalopram

We used a support vector machine (SVM) to distinguish the underlying patterns of escitalopram treatment response by classifying responders and non-responders at baseline. For the psilocybin group we could not implement a classification given the limited number of non-responder individuals. In particular, we trained five classifiers using as input the following (all with same dimensionality): global brain connectivity (GBC), in-weights of the GEC (GEC^in^), out-weights of the GEC (GEC^out^), total weights of the GEC (GEC^total^) and the perturbability map of brain regions from the FDT analysis. In each classifier, we computed one SVM for each number of features *F* in an accumulated manner (i.e., brain areas 1-*N*). In each SVM of a given number of features, for each k-fold (i.e., 1, 2, 3, …, 1000), we calculated the standard error of the mean (SSND) of brain areas between responders and non-responders in the training set, ranked them in descending order, and used the top *F* brain areas for training. We selected those same areas for the testing set.

The maximum accuracy reached in each classifier was 35.55% with 31 features for GBC, 52.95% with 5 features for GEC^in^, 58.25% with 3 features for GEC^out^, 56.85% with 69 features for GEC^total^, and 85.50% with 1 feature for FDT. Results in **Figure 5B** show the corresponding confusion matrices and statistics between classifiers. The GEC measures had higher robustness than the FC to classify subgroups of individuals. Furthermore, the deviation of the FDT presented even higher accuracy. The most relevant brain areas for all classifications were mainly from the LIM, DMN, SCN, and others from VIS, SOM and VAN (**Supplementary Table S8)**. The evolution of the accuracy for each number of features *F* in each classification can be found in **Supplementary Information Figure S7**.

### Hierarchical reconfiguration as a predictor of depression score in psilocybin

We then looked for a relation between baseline brain measures and the level of clinical improvement of the patients. We calculated the correlation between the asymmetry within the SOM, at baseline, and the change (after-before) in BDI scores (i.e., clinical depression symptoms). For psilocybin, the correlation was significantly positive, whereas for escitalopram, or when combining both groups, it was non-significant **(Figure 5C and Supplementary Information Figure S8)**. In other words, for the psilocybin arm, lower baseline asymmetry within the SOM is related to higher clinical improvements after treatment. The SOM network was chosen given it showed higher FDT deviation across time (before and after treatment) and treatment (psilocybin and escitalopram) (**Figure 2A)**, highest absolute changes between before and after treatment (**Figure 2B)**, and areas in the top differences between responders and non-responders for both treatment groups (**Figure 4A)**.

## Discussion

We successfully implemented a thermodynamic-inspired framework for analysing the hierarchy of non-equilibrium brain dynamics in patients with major depressive disorder before and after treatment with psilocybin or escitalopram **(Figure 1)**. This framework, developed by (Deco et al., 2023), proposes that asymmetrical interactions between brain areas can lead to deviations from the Fluctuation-Dissipation Theorem (FDT), characteristic of non-equilibriums systems (Kubo, 1966; Onsager, 1931). Our analysis revealed differential effects for each drug. Psilocybin significantly increased the FDT deviation after treatment compared to baseline. Furthermore, brain segregation significantly decreased, and the global, intra- and inter-module proportion of asymmetric interactions significantly increased. In other words, increased desegregation and asymmetry were found following psilocybin. We found opposite and significant results in all measures (i.e., FDT deviation, segregation and asymmetry) for the escitalopram group. These findings support and extend previous analyses in the same cohort, which suggested and showed that while both interventions reduce depressive symptoms, they do so through distinct mechanisms (Daws et al., 2022; Deco, 2024; Wall et al., 2025). Overall, by applying a thermodynamic perspective on brain function, our study offers new insights into the underlying non-equilibrium brain dynamics of depression in the post-acute effects of treatment with psilocybin and escitalopram.

The changes in FDT deviation at a whole-brain level indicate changes in the brain’s functional hierarchical organisation and distance from equilibrium dynamics in terms of the breaking of the detailed balance. Our results showed that the level of hierarchical non-equilibrium brain dynamics increased for psilocybin and decreased for escitalopram **(Figure 2A)**. Furthermore, they reflected the same trend within each therapy in the proportion of global asymmetric interactions **(Supplementary Figure S5A)**. We move a step further from the study showing global directedness decreases after psilocybin and increases after escitalopram in the same dataset (Deco, 2024). During the acute stage of psychedelic action in healthy participants, brain dynamics appear to move closer to equilibrium - based on reduced temporal asymmetry in the directionality of information flow - and higher complexity (Vohryzek, 2025). This is interpreted as a relaxation of hierarchical constraints alongside a broadened repertoire of brain substates, enabling novel and unpredictable trajectories consistent with the ‘entropic brain’ model of conscious states (Carhart-Harris, 2025). In contrast, our results showed post-acute departures from equilibrium, which we suggest, with caution, may reflect a recalibration of hierarchical organisation relative to condition-specific baseline brain dynamics in depression, compatible with clinical improvement. To further understand the different results, a future step would be to quantify these measures at the condition level (healthy versus depressed), and during the onset and peak of psychedelic action versus post-acute effects. It would also be important to carefully contextualise the underlying assumptions and nuances in the characterisation of hierarchy in each method (Nartalio-Kaluarachchi, 2025).

At a regional level, our results revealed distinct post-treatment reconfigurations in perturbability (i.e., regional FDT deviation) between psilocybin and escitalopram (**Figure 2B-C**). For both treatment groups, the perturbability of almost all areas followed their respective global trends in FDT deviation (i.e., positive for psilocybin and negative for escitalopram). The areas with highest changes in perturbability were primarily located in the SOM and VAN networks, which were also the ones orchestrating the hierarchy (i.e., highest perturbability) across sessions. Other brain regions with largest absolute differences in perturbability were located mainly in the DMN for psilocybin and DAN for escitalopram. The relevance of these networks in depression has been highlighted in several studies. In the case of the SOM, it has been associated with the severity of depression given the physical component of the disorder arising from how the body interacts with the environment (i.e., embodied phenomenon) (Ray et al., 2021). Furthermore, the salience network has been linked with cognitive vulnerability in depression (Lydon-Staley et al., 2019; Kaiser et al., 2015; Sliz & Hayley, 2012). In addition, the DMN has been hypothesised to be implicated in depression (Hamilton et al., 2015) given its association with mind-wandering, internal thoughts, self-referential thinking and rumination (Kaiser et al., 2015; Lydon-Staley et al., 2019; Mason et al., 2007; Zhou et al., 2020). Lastly, the DAN has shown group connection differences between MDD and healthy controls after escitalopram treatment (Wang et al., 2024).

We also analysed regional differences between responders and non-responders in each intervention arm. Brain area perturbability changes showed differences for each treatment associated with clinical improvements **(Figure 4A**). Regarding cortical brain areas, psilocybin responders presented increases in perturbability after treatment across almost all areas, while non-responders exhibited more heterogeneous changes with only half of the areas increasing in perturbability. On the other hand, escitalopram responders and non-responders both showed reductions in perturbability after treatment for almost all areas, with more pronounced changes for responders overall.

The segregation analysis provides insights into the global network reorganisation following each treatment. Results for the psilocybin arm showed a significant lower modularity in the GEC matrices (**Figure 3A-B)**. Interestingly, this goes in line with the acute phase of psilocybin in healthy participants characterised by global integration and communication (Roseman et al., 2014), increased/decreased connectivity between/within networks (Yu et al., 2024), and reduced functional differentiation at the extremes of the principal gradient of cortical organisation (Girn et al., 2022). Furthermore, increased global functional integration has been related with the peak (i.e., ‘mystical’) experience during psychedelic action, which is in turn associated with positive effects in mental health (Carhart-Harris, 2019). The impact of psychedelics on modularity is further discussed in (Girn et al., 2023). On the other hand, our analysis in the escitalopram arm presented a significant increase in modularity, supporting the opposite effects of the treatments. In the same cohort and timeframe as our study (i.e., MDD patients, 3 weeks and 1 day after DD2), (Daws et al., 2022) found a significant decrease in FC modularity after psilocybin therapy, which correlated with antidepressant efficacy, and no significance in neither analysis (i.e., FC modularity change and correlation with clinical scores) for the escitalopram arm. Our work extends on these previous results two-fold, revealing the opposite neural changes of psilocybin and escitalopram in a significant way, and the increased efficacy of the GEC compared to the FC to show brain changes. Lastly, using the modules obtained, we computed the within and between modular asymmetry (i.e., in terms of proportion of asymmetric interactions) and found that the global trends in FDT deviation and asymmetry are led by both inter and intra-module processing (**Figure 3C** and **Supplementary Information Figure S5B-C)**.

Furthermore, we successfully associated baseline brain measures with changes in clinical scores after treatment. For psilocybin, we found a significant positive correlation between baseline asymmetry (i.e., proportion of asymmetric interactions) within the SOM network and BDI score change (after – before) (**Figure 4C)**. In other words, lower baseline asymmetry within the SOM is related with higher clinical improvement after treatment. Interestingly, another study has already related the SOM with response to treatment of depression (Ray et al., 2021). For escitalopram, we performed a pattern separation of responders from non-responders using a SVM classifier which had as input the global brain connectivity (GBC), in-weights of the GEC (GEC^in^), out-weights of the GEC (GEC^out^), total weights of the GEC (GEC^total^) and the FDT deviation across areas. The classifier reached a maximum accuracy with the FDT deviation (85.50%), performing significantly higher than the GEC measures and the GBC (**Figure 4B)**. Different fMRI-based biomarkers of depression have been already implemented in previous studies (Pilmeyer et al., 2022), and together with our results they are interesting for the growing field of individualised treatments and precision psychiatry (Tura & Goya-Maldonado, 2023).

Our study has several limitations worth mentioning. It relies on a brain parcellation of 80 areas (i.e., DK80), which balances computational costs and spatial accuracy of the findings (Deco et al., 2021). There is no agreed-upon standard for fMRI parcellation (Eickhoff et al., 2018), but we must acknowledge that our findings are constrained to the DK80 resolution. We also used a structural connectivity template from a separate healthy cohort given we did not have individualised connectomes of the analysed cohort. Nevertheless, studies in the past have successfully implemented templates (Deco et al., 2018), as they are a neutral starting point, with relevant links in the GEC optimisation process fitted to the patient’s empirical data. Lastly, the relatively small sample size limits the generalisability of the findings. Some of the findings may not be representative of the broader population of individuals with MDD treated with psilocybin and escitalopram. Replicating the study with larger cohorts would enhance the statistical power and improve the robustness of the conclusions.

Overall, our work shows that in the context of thermodynamics, the deviation from the Fluctuation-Dissipation Theorem is a powerful tool for uncovering the hierarchical reorganisation and non-equilibrium brain dynamics accompanying interventions for major depressive disorder (Deco et al., 2023). We found significant differential effects for the interventions of psilocybin and escitalopram. Psilocybin shifts the brain towards more non-equilibrium (i.e., greater deviation from the FDT) and proportion of asymmetric interactions, and less segregation, while escitalopram produces opposite effects in all measures. To conclude, the thermodynamics framework (Kringelbach et al., 2024) is paving the way for future research aimed at optimizing existing therapeutic interventions and developing more effective treatments for depression and potentially other brain disorders.

## Data availability

The raw data can be requested to R.C.-H., the chief investigator on the original work.

## Code availability

Whole-brain models and analysis was performed in MATLAB R2022a and R2024a software from MathWorks (Natick, MA, USA). The code is available on https://github.com/pcdagnino/FDT. Other functions were used from the free available Brain Connectivity Toolbox (brain-connectivity-toolbox.net). Subcortical figures were built in Python V3.12.

## Acknowledgements

P.D. is supported by the Department of Research and Universities of the Government of Catalonia, Agency for Management of University and Research Grants (AGAUR), FI-SDUR programme (Grant No. 2022 FISDU 00229). I.A.P is supported by Grant PID2022-136216NB-100 funded by MICIU/AEI/ 10.13039/501100011033 and, as appropriate, by “ERDF A way of making Europe”, by “ERDF/EU”, by the “European Union” or by the “European Union NextGenerationEU/PRTR”. G.Z.L is partially funded by the Grant ERDF-Project Brain dynamics, No. CZ.02.01.01/00/22\_008/0004643. A.E. is supported by the European Union’s Horizon Europe research and innovation programme under the Marie Sklodowska-Curie Actions (ID: 101207460, NEUROCONTRA, HORIZON-MSCA-2024-PF-01-01), and by the project eBRAIN-Health - Actionable Multilevel Health Data (ID: 101058516) funded by the EU Horizon Europe. Y.S.P. is supported by the EU funded Project NEurological MEchanismS of Injury, and Sleep-like cellular dynamics (NEMESIS; ref. 101071900) funded by the EU ERC Synergy Horizon Europe, and by the Grant PID2024-162576NA-I00 funded by MICIU/AEI/10.13039/501100011033 and by “ERDF A way of making Europe”, ERDF, EU. M.L.K. is supported by the Centre for Eudaimonia and Human Flourishing (funded by the Pettit and Carlsberg Foundations) and Center for Music in the Brain (funded by the Danish National Research Foundation, DNRF117). G.D. is supported by Grant PID2022-136216NB-I00 funded by MICIU/AEI/10.13039/501100011033 and by “ERDF A way of making Europe”, “ERDF, EU”, Project NEurological MEchanismS of Injury, and Sleep-like cellular dynamics (NEMESIS) (ref. 101071900) funded by the EU ERC Synergy Horizon Europe, AGAUR research support grant (ref. 2021 SGR 00917) funded by the Department of Research and Universities of the Generalitat of Catalunya, and Grant PID2024-155136NI-I00 financed by MICIU/AEI/10.13039/501100011033/ and by “ERDF A way of making Europe”, ERDF, EU. The double-blind randomised controlled trial was funded by a private donation from the Alexander Mosley Charitable Trust, supplemented by Founders of Imperial College London’s Centre for Psychedelic Research. The research was conducted at The National Institute for Health and Care Research (NIHR) Imperial Clinical Research Facility (ICRF). We would like to thank the Clinical Data Systems team at the Imperial Clinical Trials Unit for their support. All authors affiliated with the Imperial College London Division of Psychiatry are supported by the National Institute of Health Research (NIHR) Imperial Biomedical Research Collaboration.

