## Supplementary Information for "Distinct brain responses to psilocybin and escitalopram in depression captured by the Fluctuation-Dissipation Theorem"

### Materials and Methods

- **Whole-brain modelling**

#### *Hopf bifurcation model*

Whole-brain dynamics can be described by coupling the local dynamics of different areas (i.e., *N* nodes) via a connectivity matrix *C*. Each brain area is simulated with a Stuart-Landau oscillator, which corresponds to the normal form of a supercritical Hopf bifurcation (i.e., standard model for examining shift from noisy to oscillatory dynamics) (Deco et al., 2017). This is expressed by the following equation:

$$\frac{dz_j}{dt} = (a_j + i\omega_j)z_j - |z_j|^2 z_j + \sum_{k=1}^{N} C_{jk}(z_k - z_j) + \eta_j, \tag{1}$$

where $z_j$ is the complex variable that denotes the state of a node $j$ ($z_j = x_j + iy_j$), $a_j$ is the bifurcation parameter, $\omega_j$ is the node frequency, and $\eta_j$ is Gaussian noise (with variance $\sigma^2$). The bifurcation parameter governs the dynamics, for $a_j > 0$ the local dynamics settle into a stable limit cycle generating self-sustained oscillations with frequency $f_j = w_j/2\pi$. On the other hand, for $a_j < 0$, the local dynamics represent a stable spiral point generating noisy oscillations if noise is present or damped oscillations otherwise. We used a value just below zero ($a_j = -0.02$) for all brain areas as it generates fluctuating stochastically structured signals that best preserve dynamically responding brain networks and resting-state network structure (Sanz Perl et al., 2023). Lastly, we calculated the intrinsic node frequency $\omega_j$ by band-pass filtering the fMRI signals (i.e., 0.008–0.08 Hz) and computing the averaged peak frequency of each brain area across all individuals. The fMRI signals are modelled by the real part of the state variables (i.e., $x_j = Real(z_j)$).

#### *Hopf linear approximation*

The finding on the best working point for fitting whole-brain models at the brink of the bifurcation (i.e., $a_j$ slightly negative and very near zero) (Sanz Perl et al., 2023) allows a linearization of the dynamics and in consequence permits an analytical solution for the functional connectivity matrix $FC$ (i.e., Pearson correlations between all pairs of brain regions) (Ponce-Alvarez & Deco, 2024). The whole-brain functional correlations can be obtained by implementing a linear noise approximation and the dynamical system in **Equation 1** can be expressed in vector form:

$$\frac{dz}{dt} = (a - S + i\omega) \odot z - (z \odot \bar{z})z + Cz + \eta. \tag{2}$$

In this equation, $z = [z_1, \ldots, z_N]^T$, $a = [a_1, \ldots, a_N]^T$, $\omega = [\omega_1, \ldots, \omega_N]^T$ and $\eta = [\eta_1, \ldots, \eta_N]^T$. In addition, $S = [S_1, \ldots, S_N]^T$ is a vector indicating the connectivity strength of each node (i.e., $S_i = \sum_j C_{ij}$). Furthermore, $[\ ]^T$ corresponds to the transpose operation, the symbol $\odot$ represents the element-wise product, and $\bar{z}$ is the complex conjugate of $z$.

The equation describes the linear fluctuations around the fixed point $z = 0$, the solution of $\frac{dz}{dt} = 0$. If the real and imaginary parts of the state variables are separated, and higher-order terms discarded (i.e., terms $(z \odot \bar{z})z$), the evolution of the linear fluctuations follow a Langevin stochastic linear equation:

$$\frac{d}{dt}\delta u = J\delta u + \eta. \quad (3)$$

Here, $\delta u = [\delta x, \delta y]^{\mathrm{T}} = [\delta x_1, \ldots, \delta x_N, \delta y_1, \ldots, \delta y_N]^{\mathrm{T}}$ is a 2N-dimensional vector which contains fluctuations of the real and imaginary state variables. Furthermore, $J$ is a $2N\ X\ 2N$ Jacobian matrix of the system evaluated at the fixed point, with the following form as block matrix:

$$J = \begin{bmatrix} Jxx & Jxy \\ Jyx & Jyy \end{bmatrix}. \quad (4)$$

The matrices $J_{xx}$ , $J_{xy}$, $J_{yx}$ and $J_{yy}$ have a size of $N\ X\ N$; $J_{xx} = J_{yy} = diag(a - S) + C$ and $J_{xy} = -J_{yx} = diag(\omega)$ , in which $diag(v)$ corresponds to the diagonal matrix with the vector $v$ as the diagonal. As a note, the linearization is possible only if $z = 0$ is a stable solution of the system, which corresponds to $J$ having all eigenvalues with a negative real part.

To calculate the covariance matrix $K = \langle \delta u \delta u^T \rangle$, **Equation 3** is expressed as $d\delta u = J\delta u dt + dW$. Here, $dW$is a 2N-dimensional Wiener process, whose covariance is $\langle dWdW^T \rangle = Qdt$, and $Q$ is the noise covariance matrix (which is diagonal if the noise is uncorrelated). Then, using Itô's stochastic calculus, $d(\delta u \delta u^T) = d(\delta u)\delta u^T + \delta u d(\delta u^T) + d(\delta u)d(\delta u^T)$. Taking expectations, noting $\langle \delta u dW^T \rangle = 0$, and keeping terms to first order in the differential $dt$, the following equation is obtained:

$$\frac{dK}{dt} = JK + KJ^T + Q. \quad (5)$$

Solving this equation for the case $\frac{dK}{dt} = 0$, the stationary covariance can be obtained analytically using the eigen-decomposition of the Jacobian matrix $J$ (Deco et al., 2023).

***Model optimisation: Generative Effective Connectivity***

The coupling connectivity matrix $C$ is optimised with a pseudo-gradient procedure until the best fit between the model and the empirical data is achieved. We used as a starting point for the optimisation process a standard structural connectivity matrix from the diffusion MRI data of the Human Connectome Project (Deco et al., 2021). We performed 1500 iterations updating each known existing pair of anatomical connections, except for homotopic areas (i.e., corresponding brain areas in opposite hemispheres), which should be also updated given these are captured less accurately in tractography. In each iteration, the corresponding connections were updated by adding the difference between the simulated and the empirical functional correlation matrix and normalised time-shifted covariance matrix:

$$C_{ij} = C_{ij} + \alpha(FC_{ij}^{emp} - FC_{ij}^{sim}) + \varsigma(FS_{ij}^{emp}(\tau) - FS_{ij}^{sim}(\tau)). \quad (6)$$

Here, $\alpha$ and $\varsigma$ were set to $0.0004$ and $0.0001$, respectively. The empirical functional connectivity ($FC^{emp}$) is obtained by computing the normalised covariance matrix of the timeseries. In addition, the empirical normalised time-shifted covariance ($FS^{emp}(\tau)$) is calculated by dividing each pair of regions *(i,j)* from the shifted covariance matrix ($KS^{emp}(\tau)$) by $\sqrt{KS_{ii}^{emp}(0)KS_{jj}^{emp}(0)}$. Furthermore, the simulated functional connectivity ($FC^{sim}$) is generated with the first $N$ rows and columns of the simulated covariance matrix $K$, which is the real part of the dynamics and represents the BOLD fMRI signal. Lastly, the simulated normalised time-shifted covariance ($FS^{sim}(\tau)$) is obtained after selecting the first *N* rows and columns of the simulated time-shifted covariance ($KS^{sim}(\tau)$), and dividing each pair of regions ($i$,$j$) by $\sqrt{KS_{ii}^{sim}(0)KS_{jj}^{sim}(0)}$. Here,

$$KS^{sim}(\tau) = exp(\tau\, J)K, \tag{7}$$

where $KS^{sim}(0) = K$.

Every 100 iterations, if the difference between empirical and modelled measures was higher than the difference from the previous iteration, or smaller than 0.001 from the last iteration, the procedure broke and the output matrix from this iteration was used. The final optimised matrix C corresponds to the Generative Effective Connectivity (GEC) matrix. The GEC is asymmetric given the optimisation includes time-shifted matrices which break the symmetry (time-shift set to $\tau = 2$), leading to non-equilibrium dynamics therefore allowing for an assessment of the violation of the FDT (Kringelbach et al., 2023). We performed the optimisation pipeline in a two-step manner, first calculating the GEC at a group level (psilocybin baseline, psilocybin after treatment, escitalopram baseline, escitalopram after treatment) by using the DTI as a starting point. Then, we calculated the GEC at an individual level, using the GEC of the patient's group as a starting point.

**Figures**

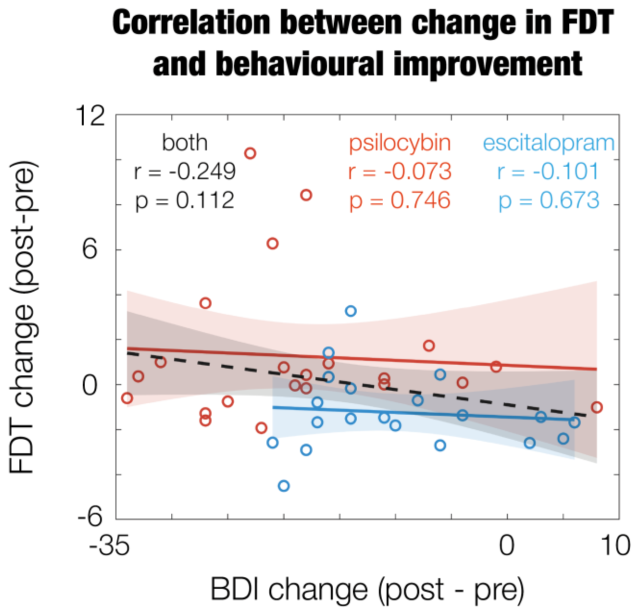


***Supplementary Figure S1: Correlation between change in FDT deviation and BDI scores.*** *Psilocybin in red, escitalopram in blue, both groups combined in black. The change in FDT deviation does not correlate significantly with the change in BDI scores for either group separately nor when combined.*

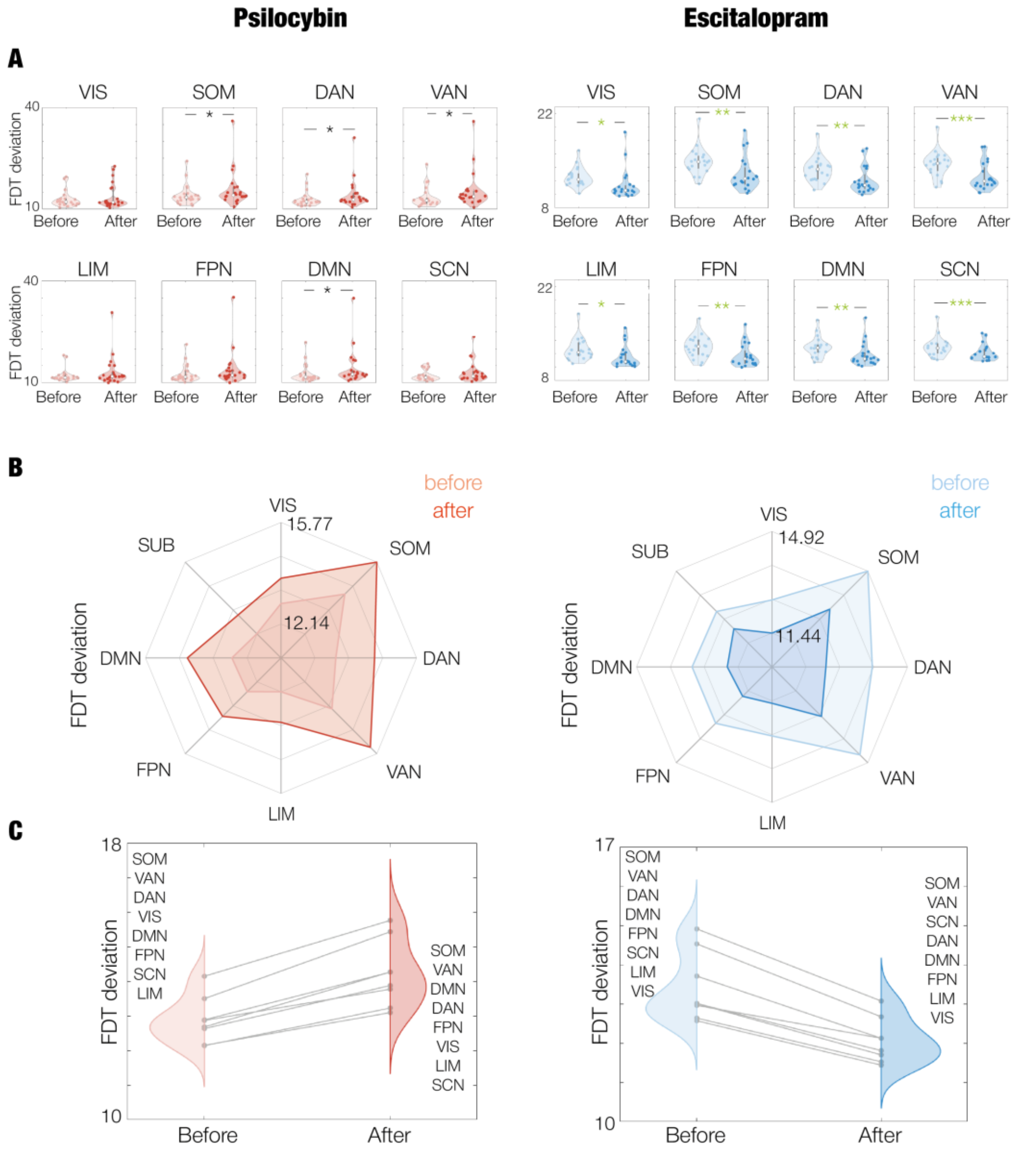


***Supplementary Figure S2: FDT deviation values across resting-state networks following administration of psilocybin and escitalopram.*** *Psilocybin and escitalopram treatments can be observed in the left and right columns, respectively.* ***A.*** *Repeated measures plot for the FDT deviation for each resting-state network. In a plot, each point is the weighted average FDT deviation for all brain areas belonging to the specific network under evaluation for each individual. The white middle dot of the violin corresponds to the median of all individuals. There are significant differences between baseline and after treatment for several networks in both psilocybin and escitalopram groups. One asterisk (i.e.,*) represents $p < 0.05$, two asterisks (i.e., **) represents $p < 0.01$. The green asterisks are the ones that remain significant after correction by multiple comparisons with FDR. The trend is opposite for each treatment. In the psilocybin arm, none survive correction by multiple comparisons. For escitalopram, all survive correction by multiple comparisons.* ***B.*** *Spider plots show the mean FDT deviation of the resting-state networks before and after each intervention.* ***C****. Repeated measures plot showing the ranking of each resting-state network before and after treatment.*

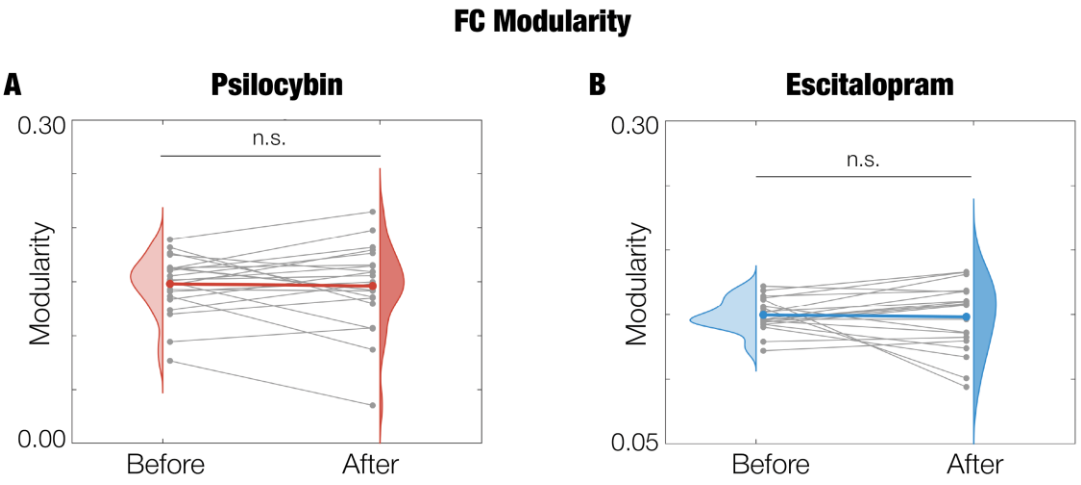


***Supplementary Figure S3: Modularity of Functional Connectivity.*** *We measured brain network segregation by computing the modularity with Louvain algorithm and Newman as cost-function. Grey lines represent the trajectory of each individual whereas the coloured line represents their averaged trajectory within each intervention arm. There are no significant differences (n.s.) for either* ***A.*** *Psilocybin group nor* ***B.*** *Escitalopram group.*

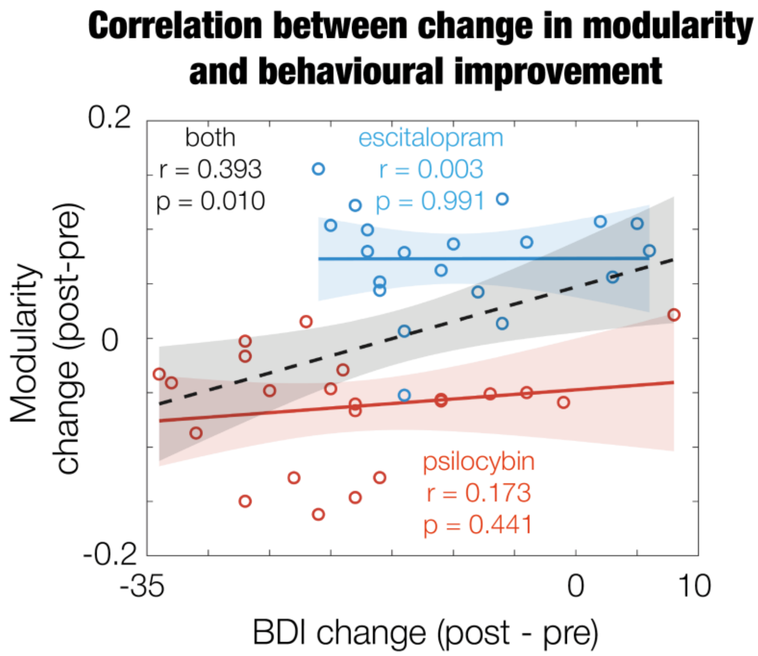


***Supplementary Figure S4: Correlation between change in modularity and BDI scores.*** *Psilocybin in orange, escitalopram in blue, both groups combined in black. We correlated the change in modularity with the change in BDI. We only found significance when combining both groups (psilocybin and escitalopram).*

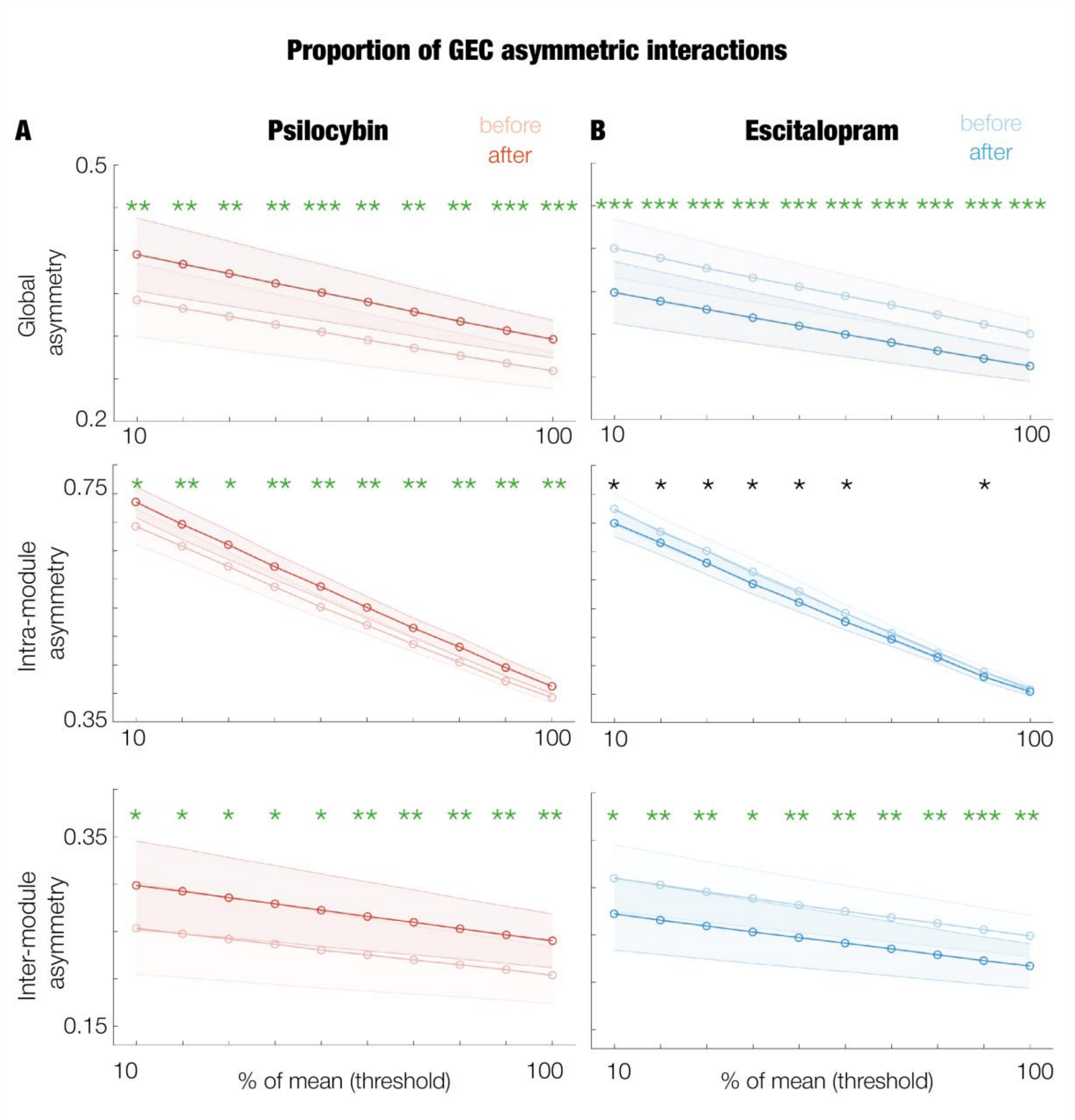


***Supplementary Figure S5. Asymmetry analysis of the GEC at a subject level for psilocybin and escitalopram.*** *We quantified asymmetric interactions as the proportion of cells that remain asymmetric in a thresholded absolute matrix minus its transpose. Results across thresholds are shown for psilocybin (left column) and escitalopram (right column) for* ***A.*** *Global GEC,* ***B.*** *Intra-module GEC and* ***C.*** *Inter-module GEC. Modules were obtained using the Louvain algorithm and Newman cost-function. Significance between before and after asymmetry for each group is represented by asterisks (*, p < 0.05; **, p < 0.01; and ***, p < 0.001). Green asterisks correspond to survivors after correction by multiple comparisons.*

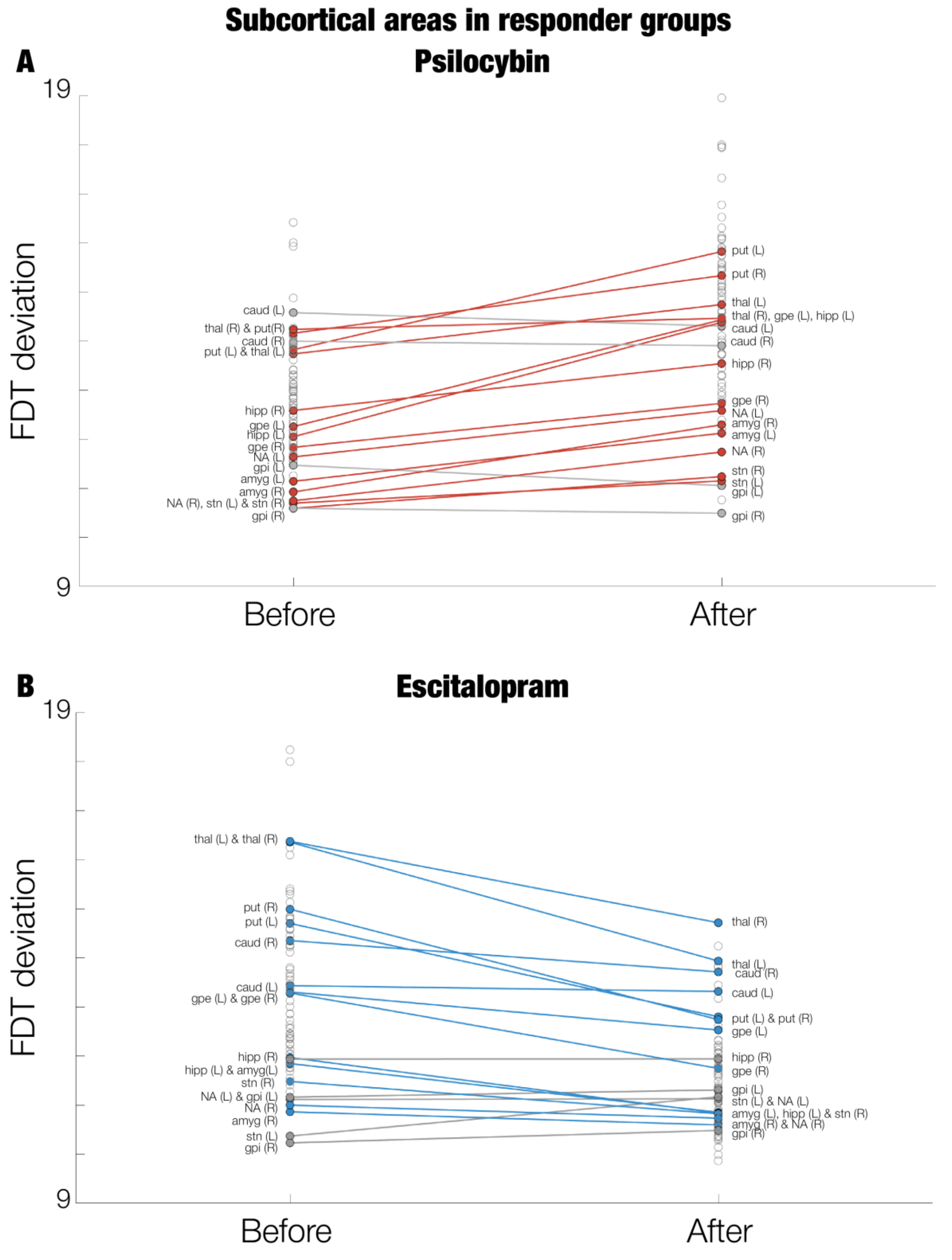


***Supplementary Figure S6: Subcortical areas in terms of their ranking in FDT deviation.*** *Each point corresponds to a brain area averaged across participants for before and after treatment in **(A)** Psilocybin and **(B)** Escitalopram**.** Subcortical areas are coloured, labelled, and a line connecting each area from before and after treatment is drawn. In colour are the brain areas following the global trends of the FDT deviation, positive change for psilocybin in red and negative change for escitalopram in blue.*

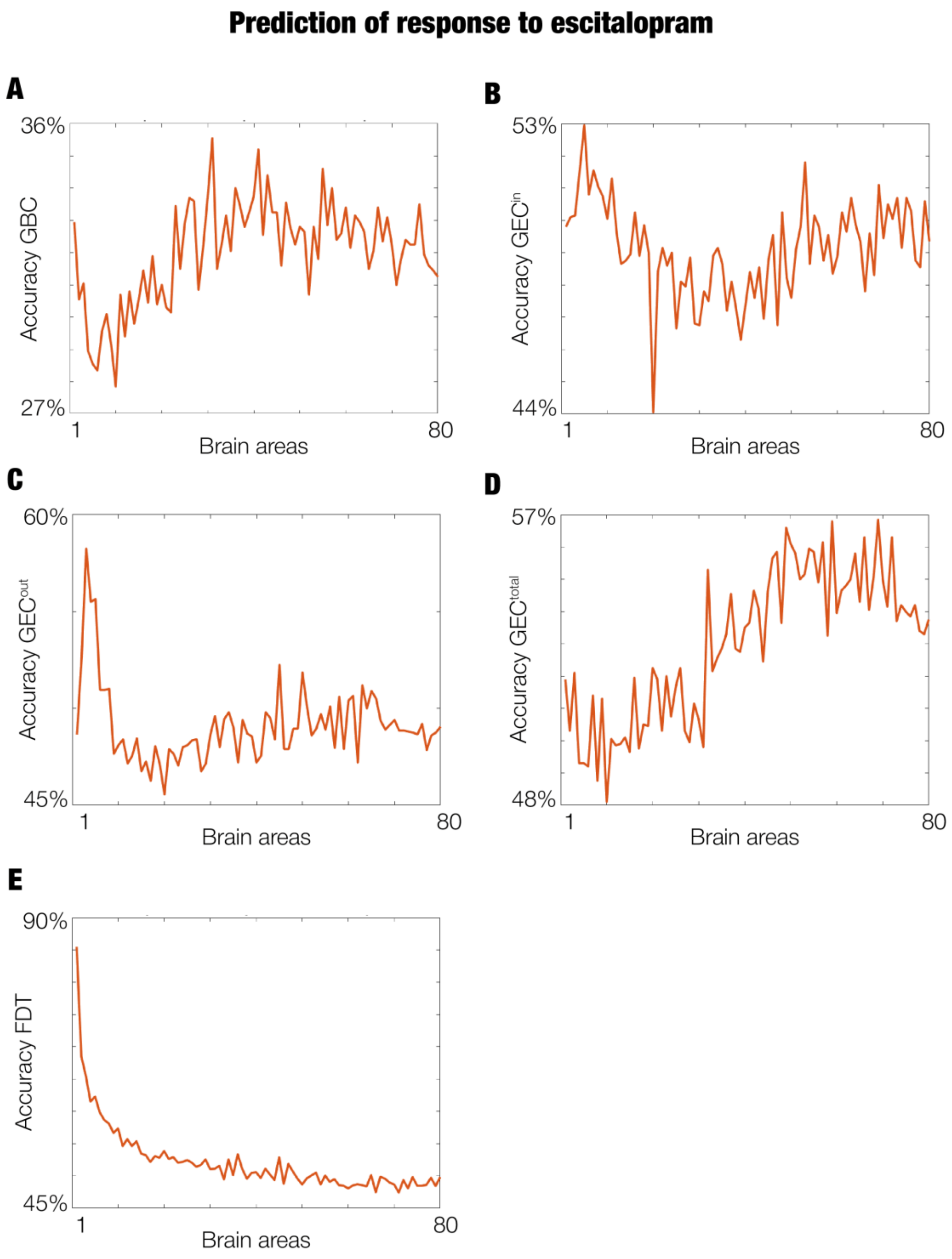


***Supplementary Figure S7: SVM to predict response to escitalopram treatment.*** *We trained a SVM classifier between baseline responders and non-responders in the escitalopram group. We studied the predictive power of brain areas by sequentially adding one by one in a cumulative manner. For each number of features, for each k-fold, we split the data into train and test sets. In each fold, with the train set, we calculated the standard error of the mean (SSND) of brain areas between responders and non-responders, and ranked the brain areas in descending order based on their SSND values. Then, we used the corresponding number of features with the highest SSND. We implemented the classifier using as input (**A**) GBC, (**B**) $GEC^{in}$ , (**C**) $GEC^{out}$ , (**D**) $GEC^{total}$ , and (**E**) FDT.*

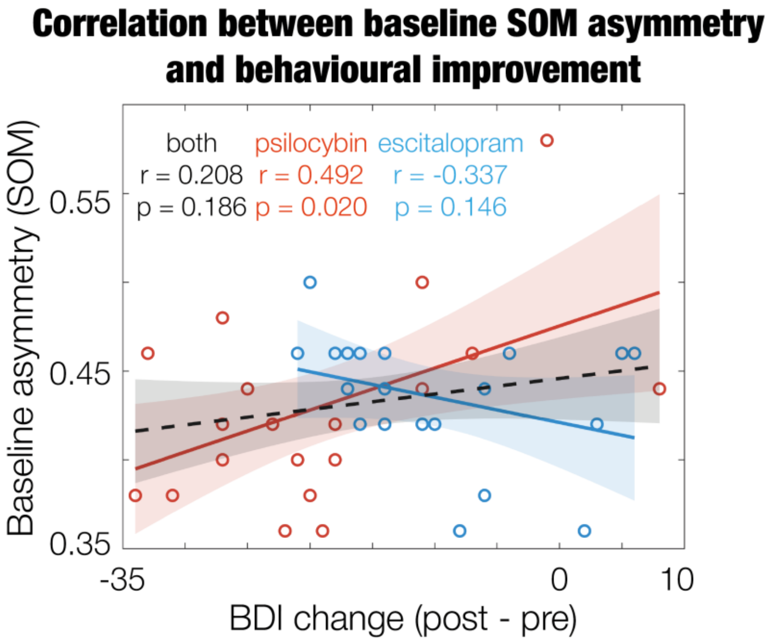


***Supplementary Figure S8: Correlation between baseline asymmetry within the SOM and the change in BDI scores.*** *We implemented the mean as threshold for the asymmetry computation to balance filtered noise and meaningful asymmetry patterns preservation. Psilocybin in orange, escitalopram in blue, both groups combined in black. The baseline asymmetry within the SOM correlates significantly with the change in BDI for psilocybin but not for escitalopram nor when combining both groups.*

Y: FDT
Factor 1: Binary classification for responders and non-responders
Factor 2: BDI change (post-pre)
Factor 3: Treatment (psylocibin or escitalopram)

N-way anova, constrained (Type III) sums of squares.

Y ~ 1 + Factor1 + Factor2 + Factor3

| Source | Sum Sq. | d.f. | Mean Sq. | F | Prob>F |
|---|---|---|---|---|---|
| Factor1 | 20.119 | 1 | 20.1189 | 3.02 | 0.1043 |
| Factor2 | 173.423 | 25 | 6.9369 | 1.04 | 0.4849 |
| Factor3 | 66.717 | 1 | 66.7169 | 10.01 | 0.0069 |
| Error | 93.353 | 14 | 6.6681 | | |
| Total | 332.131 | 41 | | | |

***Supplementary Table S1: Analysis of variance (ANOVA) with different variables.*** *Treatment is the only significant factor.*

| | Psilocybin | | Escitalopram | |
|---|---|---|---|---|
| Top 20% | right posterior cingulate | VentralAttn | left superior temporal | Somatomotor |
| | left superior temporal | Somatomotor | right superior temporal | Somatomotor |
| | right superior temporal | Somatomotor | left posterior cingulate | VentralAttn |
| | left posterior cingulate | VentralAttn | right thalamus | Subcortical |
| | right superior frontal | Default | right posterior cingulate | VentralAttn |
| | left transverse temporal | Somatomotor | left thalamus | Subcortical |
| | left postcentral | Somatomotor | left insula | VentralAttn |
| | left caudate | Subcortical | left precentral | Somatomotor |
| | right precentral | Somatomotor | left supramarginal | VentralAttn |
| | left precentral | Somatomotor | right precentral | Somatomotor |
| | right paracentral | Somatomotor | right insula | VentralAttn |
| | right postcentral | Somatomotor | right putamen | Subcortical |
| | right transverse temporal | Somatomotor | left caudal anterior cingulate | VentralAttn |
| | left paracentral | Somatomotor | left postcentral | Somatomotor |
| | right caudate | Subcortical | right superior frontal | Default |
| | left pericalcarine | Visual | right supramarginal | VentralAttn |
| Bottom 20% | right pars orbitalis | Default | right lateral orbitofrontal | Limbic |
| | left medial orbitofrontal | Limbic | right parahippocampal | Visual |
| | left pars triangularis | Default | right pars orbitalis | Default |
| | right lateral orbitofrontal | Limbic | right caudal middle frontal | Frontoparietal |
| | left lateral orbitofrontal | Limbic | left stn | Subcortical |
| | left gpi | Subcortical | left gpi | Subcortical |
| | left nucleus accumbens | Subcortical | right rostral middle frontal | Frontoparietal |
| | left pars orbitalis | Default | right lateral occipital | Visual |
| | left parahippocampal | Visual | right medial orbitofrontal | Limbic |
| | left amygdala | Subcortical | left pars orbitalis | Default |
| | left entorhinal | Limbic | left parahippocampal | Visual |
| | right amygdala | Subcortical | left medial orbitofrontal | Limbic |
| | right gpi | Subcortical | right nucleus accumbens | Subcortical |
| | right stn | Subcortical | right amygdala | Subcortical |
| | right nucleus accumbens | Subcortical | left nucleus accumbens | Subcortical |
| | left stn | Subcortical | right gpi | Subcortical |

***Supplementary Table S2: Areas at the top and bottom of the hierarchy ordered by FDT deviation before treatment.*** *The table shows the top (first row) and bottom (second row) 20% of brain areas – and their corresponding RSN – in descending order of FDT deviation. Before treatment for the psilocybin (first column) and escitalopram (second column) groups for all patients.*

| | Psilocybin | | Escitalopram | |
|---|---|---|---|---|
| Top 20% | left superior temporal | Somatomotor | left posterior cingulate | VentralAttn |
| | right superior temporal | Somatomotor | left superior temporal | Somatomotor |
| | left posterior cingulate | VentralAttn | right posterior cingulate | VentralAttn |
| | right posterior cingulate | VentralAttn | right thalamus | Subcortical |
| | left insula | VentralAttn | right superior temporal | Somatomotor |
| | right middle temporal | Default | left thalamus | Subcortical |
| | left precentral | Somatomotor | right caudate | Subcortical |
| | left transverse temporal | Somatomotor | left caudate | Subcortical |
| | right superior frontal | Default | left precentral | Somatomotor |
| | left caudal anterior cingulate | VentralAttn | left postcentral | Somatomotor |
| | right insula | VentralAttn | left putamen | Subcortical |
| | left postcentral | Somatomotor | right middle temporal | Default |
| | right precentral | Somatomotor | left transverse temporal | Somatomotor |
| | left paracentral | Somatomotor | right precentral | Somatomotor |
| | left putamen | Subcortical | right putamen | Subcortical |
| | right caudal anterior cingulate | VentralAttn | right transverse temporal | Somatomotor |
| Bottom 20% | left pars orbitalis | Default | right amygdala | Subcortical |
| | left nucleus accumbens | Subcortical | left rostral anterior cingulate | Default |
| | left caudal middle frontal | Default | left medial orbitofrontal | Limbic |
| | left rostral middle frontal | Frontoparietal | right medial orbitofrontal | Limbic |
| | right amygdala | Subcortical | right pars orbitalis | Default |
| | left medial orbitofrontal | Limbic | left rostral middle frontal | Frontoparietal |
| | right medial orbitofrontal | Limbic | right rostral middle frontal | Frontoparietal |
| | left entorhinal | Limbic | left nucleus accumbens | Subcortical |
| | left amygdala | Subcortical | left entorhinal | Limbic |
| | right rostral middle frontal | Frontoparietal | left caudal middle frontal | Default |
| | right nucleus accumbens | Subcortical | right nucleus accumbens | Subcortical |
| | right stn | Subcortical | right gpi | Subcortical |
| | left stn | Subcortical | right caudal middle frontal | Frontoparietal |
| | left gpi | Subcortical | left pars orbitalis | Default |
| | right entorhinal | Limbic | left pars triangularis | Default |
| | right gpi | Subcortical | left parahippocampal | Visual |

***Supplementary Table S3: Areas at the top and bottom of the hierarchy ordered by FDT deviation after treatment.*** *The table shows the top (first row) and bottom (second row) 20% of brain areas – and their corresponding RSN - in descending order of FDT deviation. After treatment for the psilocybin (first column) and escitalopram (second column) groups for all patients.*

| | Psilocybin | | | Escitalopram | | |
|---|---|---|---|---|---|---|
| Top 20% change | left insula | VentralAttn | 3.275 | right hippocampus | Subcortical | 0.431 |
| | right middle temporal | Default | 2.704 | | | |
| | left supramarginal | VentralAttn | 2.518 | | | |
| | left caudal anterior cingulate | VentralAttn | 2.502 | | | |
| | left superior temporal | Somatomotor | 2.477 | | | |
| | left precentral | Somatomotor | 2.120 | | | |
| | left putamen | Subcortical | 2.060 | | | |
| | right supramarginal | VentralAttn | 2.009 | | | |
| | right rostral anterior cingulate | Default | 2.007 | | | |
| | right caudal anterior cingulate | VentralAttn | 1.993 | | | |
| | right insula | VentralAttn | 1.992 | | | |
| | left lateral orbitofrontal | Limbic | 1.960 | | | |
| | left parahippocampal | Visual | 1.910 | | | |
| | right superior temporal | Somatomotor | 1.893 | | | |
| | left middle temporal | Default | 1.855 | | | |
| | right inferior temporal | Limbic | 1.838 | | | |
| Bottom 20% change | left pericalcarine | Visual | -0.018 | right superior frontal | Default | -1.890 |
| | left caudate | Subcortical | -0.074 | left superior frontal | Default | -1.925 |
| | right caudate | Subcortical | -0.095 | right putamen | Subcortical | -1.938 |
| | right gpi | Subcortical | -0.172 | left thalamus | Subcortical | -1.979 |
| | left gpi | Subcortical | -0.563 | left cuneus | Visual | -1.990 |
| | right entorhinal | Limbic | -1.127 | right caudal anterior cingulate | VentralAttn | -1.992 |
| | | | | right precentral | Somatomotor | -2.009 |
| | | | | left superior parietal | DorsalAttn | -2.048 |
| | | | | left caudal anterior cingulate | VentralAttn | -2.082 |
| | | | | right superior parietal | DorsalAttn | -2.179 |
| | | | | right supramarginal | VentralAttn | -2.263 |
| | | | | right insula | VentralAttn | -2.317 |
| | | | | left supramarginal | VentralAttn | -2.360 |
| | | | | left insula | VentralAttn | -2.872 |
| | | | | left superior temporal | Somatomotor | -3.218 |
| | | | | right superior temporal | Somatomotor | -3.489 |

***Supplementary Table S4: Changes in FDT deviation with treatment.*** *The table shows the top 20% of brain areas – and their corresponding RSN and changes in FDT deviation - with the largest positive (first row) and negative (second row) change in FDT deviation in descending order. For psilocybin (first column) and escitalopram (second column) treatment, for all patients.*

| Psilocybin | | |
|---|---|---|
| left parahippocampal | Visual | **<0.001** |
| left insula | VentralAttn | **0.001** |
| right amygdala | Subcortical | 0.003 |
| left supramarginal | VentralAttn | 0.003 |
| right middle temporal | Default | 0.004 |
| left lateral orbitofrontal | Limbic | 0.011 |
| right stn | Subcortical | 0.011 |
| left precentral | Somatomotor | 0.016 |
| right inferior temporal | Limbic | 0.016 |
| right parahippocampal | Visual | 0.017 |
| right inferior parietal | Default | 0.017 |
| left caudal anterior cingulate | VentralAttn | 0.017 |
| right supramarginal | VentralAttn | 0.018 |
| right precuneus | Default | 0.023 |
| left putamen | Subcortical | 0.023 |
| left superior temporal | Somatomotor | 0.024 |
| right pars opercularis | VentralAttn | 0.025 |
| right pars triangularis | Frontoparietal | 0.026 |
| left superior frontal | Default | 0.030 |
| right insula | VentralAttn | 0.030 |
| right rostral anterior cingulate | Default | 0.034 |
| left middle temporal | Default | 0.035 |
| left fusiform | Visual | 0.035 |
| left inferior parietal | Default | 0.038 |
| left rostral anterior cingulate | Default | 0.039 |
| left precuneus | Default | 0.039 |
| right fusiform | Visual | 0.041 |
| right superior temporal | Somatomotor | 0.042 |

| Escitalopram | | |
|---|---|---|
| left insula | VentralAttn | **<0.001** |
| right supramarginal | VentralAttn | **<0.001** |
| left supramarginal | VentralAttn | **<0.001** |
| right superior parietal | DorsalAttn | **<0.001** |
| right insula | VentralAttn | **<0.001** |
| right caudal anterior cingulate | VentralAttn | **0.001** |
| right superior temporal | Somatomotor | **0.001** |
| right putamen | Subcortical | **0.001** |
| left caudal anterior cingulate | VentralAttn | **0.002** |
| left superior temporal | Somatomotor | **0.002** |
| right gpe | Subcortical | **0.003** |
| left superior parietal | DorsalAttn | **0.004** |
| right precentral | Somatomotor | **0.005** |
| left thalamus | Subcortical | **0.005** |
| left caudal middle frontal | Default | **0.006** |
| right postcentral | Somatomotor | **0.006** |
| left lateral orbitofrontal | Limbic | **0.006** |
| right pars opercularis | VentralAttn | **0.009** |
| right inferior parietal | Default | **0.009** |
| left precentral | Somatomotor | **0.009** |
| left pars triangularis | Default | **0.009** |
| left postcentral | Somatomotor | **0.009** |
| right rostral anterior cingulate | Default | **0.010** |
| left cuneus | Visual | **0.012** |
| right pars triangularis | Frontoparietal | **0.012** |
| right caudal middle frontal | Frontoparietal | **0.013** |
| left superior frontal | Default | **0.013** |
| left pars opercularis | VentralAttn | **0.014** |
| left lingual | Visual | **0.014** |
| right superior frontal | Default | **0.015** |
| right thalamus | Subcortical | **0.015** |
| left transverse temporal | Somatomotor | **0.015** |
| right transverse temporal | Somatomotor | **0.016** |
| right fusiform | Visual | **0.020** |
| left gpe | Subcortical | **0.022** |
| right posterior cingulate | VentralAttn | **0.023** |
| left entorhinal | Limbic | **0.023** |
| left rostral anterior cingulate | Default | 0.028 |
| right paracentral | Somatomotor | 0.028 |
| left pericalcarine | Visual | 0.030 |
| left rostral middle frontal | Frontoparietal | 0.030 |
| left hippocampus | Subcortical | 0.031 |
| left parahippocampal | Visual | 0.032 |
| left paracentral | Somatomotor | 0.033 |
| right pars orbitalis | Default | 0.033 |
| right middle temporal | Default | 0.034 |
| left inferior parietal | Default | 0.036 |
| right cuneus | Visual | 0.040 |
| left pars orbitalis | Default | 0.041 |
| left amygdala | Subcortical | 0.044 |

| | | | |
|---|---|---|---|
| | left putamen | Subcortical | 0.045 |
| | right rostral middle frontal | Frontoparietal | 0.048 |
| | right inferior temporal | Limbic | 0.049 |

***Supplementary Table S5: Significant changes in FDT deviation between before and after treatment.*** *The table shows the significant brain areas in FDT deviation between before and after treatment – and their corresponding RSN and p-values - for psilocybin (first column) and escitalopram (second column) treatment, for all patients. The ones surviving correction after multiple comparisons are in bold.*

| | Psilocybin responders | | | Escitalopram responders | | |
|---|---|---|---|---|---|---|
| Top 20% change | left insula | VentralAttn | 3.399 | left stn | Subcortical | 0.798 |
| | left superior temporal | Somatomotor | 2.951 | right gpi | Subcortical | 0.254 |
| | right middle temporal | Default | 2.932 | left gpi | Subcortical | 0.148 |
| | left supramarginal | VentralAttn | 2.501 | left nucleus accumbens | Subcortical | 0.015 |
| | left lateral orbitofrontal | Limbic | 2.474 | right hippocampus | Subcortical | 0.003 |
| | left precentral | Somatomotor | 2.422 | | | |
| | left caudal anterior cingulate | VentralAttn | 2.386 | | | |
| | left hippocampus | Subcortical | 2.330 | | | |
| | left isthmus cingulate | Default | 2.294 | | | |
| | left fusiform | Visual | 2.280 | | | |
| | right insula | VentralAttn | 2.268 | | | |
| | right rostral anterior cingulate | Default | 2.261 | | | |
| | right inferior parietal | Default | 2.241 | | | |
| | right supramarginal | VentralAttn | 2.206 | | | |
| | left gpe | Subcortical | 2.177 | | | |
| | left transverse temporal | Somatomotor | 2.169 | | | |
| Bottom 20% change | right caudate | Subcortical | -0.091 | left cuneus | Visual | -2.639 |
| | right gpi | Subcortical | -0.100 | right posterior cingulate | VentralAttn | -2.650 |
| | left caudate | Subcortical | -0.275 | right fusiform | Visual | -2.661 |
| | left gpi | Subcortical | -0.415 | right precentral | Somatomotor | -2.692 |
| | right entorhinal | Limbic | -1.423 | left superior frontal | Default | -2.711 |
| | | | | left postcentral | Somatomotor | -2.764 |
| | | | | left paracentral | Somatomotor | -2.796 |
| | | | | right superior frontal | Default | -2.807 |
| | | | | left superior parietal | DorsalAttn | -2.989 |
| | | | | right supramarginal | VentralAttn | -2.997 |
| | | | | right superior parietal | DorsalAttn | -3.032 |
| | | | | left precentral | Somatomotor | -3.048 |
| | | | | left insula | VentralAttn | -3.278 |
| | | | | left supramarginal | VentralAttn | -3.400 |
| | | | | left superior temporal | Somatomotor | -3.999 |
| | | | | right superior temporal | Somatomotor | -4.189 |

***Supplementary Table S6: Changes in FDT deviation in responder groups within each treatment.*** *The table shows the top 20% of brain areas – and their corresponding RSN and changes in FDT deviation - with the largest positive (first row) and negative (second row) changes in FDT deviation in descending order. Left and right columns show the responder groups of psilocybin and escitalopram, respectively.*

| | Psilocybin non-responders | | | Escitalopram non-responders | | |
|---|---|---|---|---|---|---|
| Top 20% change | right thalamus | Subcortical | 3.075 | right hippocampus | Subcortical | 0.716 |
| | left caudal anterior cingulate | VentralAttn | 3.025 | right parahippocampal | Visual | 0.560 |
| | right caudal anterior cingulate | VentralAttn | 2.882 | left caudate | Subcortical | 0.076 |
| | left insula | VentralAttn | 2.718 | right caudate | Subcortical | 0.035 |
| | left supramarginal | VentralAttn | 2.593 | | | |
| | right amygdala | Subcortical | 2.516 | | | |
| | left parahippocampal | Visual | 2.350 | | | |
| | left putamen | Subcortical | 2.334 | | | |
| | left posterior cingulate | VentralAttn | 2.329 | | | |
| | right pars triangularis | Frontoparietal | 2.218 | | | |
| | right inferior temporal | Limbic | 1.919 | | | |
| | left pars opercularis | VentralAttn | 1.898 | | | |
| | left pars triangularis | Default | 1.715 | | | |
| | left nucleus accumbens | Subcortical | 1.705 | | | |
| | right middle temporal | Default | 1.677 | | | |
| | left thalamus | Subcortical | 1.440 | | | |
| Bottom 20% change | left isthmus cingulate | Default | -0.679 | left lingual | Visual | -1.481 |
| | left inferior parietal | Default | -0.700 | right precentral | Somatomotor | -1.555 |
| | left cuneus | Visual | -0.777 | left cuneus | Visual | -1.557 |
| | left lateral occipital | Visual | -0.811 | right superior parietal | DorsalAttn | -1.609 |
| | right paracentral | Somatomotor | -1.031 | left supramarginal | VentralAttn | -1.666 |
| | right lingual | Visual | -1.036 | left thalamus | Subcortical | -1.683 |
| | left hippocampus | Subcortical | -1.153 | left caudal anterior cingulate | VentralAttn | -1.716 |
| | right cuneus | Visual | -1.170 | right putamen | Subcortical | -1.730 |
| | left gpi | Subcortical | -1.228 | right supramarginal | VentralAttn | -1.774 |
| | left medial orbitofrontal | Limbic | -1.249 | right caudal anterior cingulate | VentralAttn | -1.836 |
| | right transverse temporal | Somatomotor | -1.300 | left entorhinal | Limbic | -1.879 |
| | right medial orbitofrontal | Limbic | -1.324 | right thalamus | Subcortical | -1.899 |
| | right lateral occipital | Visual | -1.357 | right insula | VentralAttn | -2.176 |
| | left lingual | Visual | -1.371 | left insula | VentralAttn | -2.601 |
| | left pericalcarine | Visual | -1.751 | left superior temporal | Somatomotor | -2.698 |
| | right pericalcarine | Visual | -1.972 | right superior temporal | Somatomotor | -3.023 |

***Supplementary Table S7: Changes in FDT deviation in non-responder groups within each treatment.*** *The table shows the top 20% of brain areas – and their corresponding RSN and changes in FDT deviation - with the largest positive (first row) and negative (second row) changes in FDT deviation in descending order. Left and right columns show the non-responder groups of psilocybin and escitalopram, respectively.*

| GBC | $GEC^{in}$ | $GEC^{out}$ | $GEC^{total}$ | FDT |
|---|---|---|---|---|
| right inferior temporal (Limbic) | left gpi (Subcortical) | left gpi (Subcortical) | right lingual (Visual) | left stn (Subcortical) |
| right transverse temporal (Somatomotor) | left stn (Subcortical) | right inferior temporal (Limbic) | right inferior temporal (Limbic) | left gpi (Subcortical) |
| right middle temporal (Default) | right inferior temporal (Limbic) | right lingual (Visual) | left gpi (Subcortical) | left pars orbitalis (Default) |
| left medial orbitofrontal (Limbic) | right lingual (Visual) | left inferior temporal (Limbic) | right superior temporal (Somatomotor) | left lateral occipital (Visual) |
| left inferior temporal (Limbic) | left inferior temporal (Limbic) | right inferior parietal (Default) | left gpe (Subcortical) | |
| left gpi (Subcortical) | left caudal middle frontal (Default) | left stn (Subcortical) | right caudal anterior cingulate (VentralAttn) | |
| right caudal anterior cingulate (VentralAttn) | right inferior parietal (Default) | left middle temporal (Default) | right pericalcarine (Visual) | |
| left rostral anterior cingulate (Default) | left pars orbitalis (Default) | left pars orbitalis (Default) | left caudal middle frontal (Default) | |
| right superior frontal (Default) | left pars triangularis (Default) | right middle temporal (Default) | left inferior temporal (Limbic) | |
| left stn (Subcortical) | right gpi (Subcortical) | left caudal middle frontal (Default) | left caudate (Subcortical) | |

***Supplementary Table S8: Top features (i.e., brain areas) for classifying response in the escitalopram group.*** *These correspond to the maximum classifications reached in each case.*
